# Quantitative lineage analysis identifies a long-term progenitor niche for the hepato-pancreato-biliary organ system

**DOI:** 10.1101/2020.08.06.240176

**Authors:** David Willnow, Uwe Benary, Anca Margineanu, Maria Lillina Vignola, Igor M. Pongrac, Zahra Karimaddini, Alessandra Vigilante, Jana Wolf, Francesca M. Spagnoli

## Abstract

Single cell-based studies have revealed tremendous cellular heterogeneity in stem cell and progenitor compartments, suggesting continuous differentiation trajectories with intermixing of cells at various states of lineage commitment and notable degree of plasticity during organogenesis^1–5^.

The hepato-pancreato-biliary organ system relies on a small endoderm progenitor compartment that gives rise to a variety of different adult tissues, including liver, pancreas, gallbladder, and extra-hepatic bile ducts^6, 7^. Experimental manipulation of various developmental signals in the mouse embryo underscored an important cellular plasticity in this embryonic territory^6, 8^. This is also reflected in the existence of human genetic syndromes as well as congenital or environmentally-caused human malformations featuring multiorgan phenotypes in liver, pancreas and gallbladder^6, 8^. Nevertheless, the precise lineage hierarchy and succession of events leading to the segregation of an endoderm progenitor compartment into hepatic, biliary, and pancreatic structures are not yet established. Here, we combine computational modelling approaches with genetic lineage tracing to assess the tissue dynamics accompanying the ontogeny of the hepato-pancreato-biliary organ system. We show that a long-term multipotent progenitor domain persists at the border between liver and pancreas, even after pancreatic fate is specified, contributing to the formation of several organ derivatives, including the liver. Moreover, using single-cell RNA sequencing we define a specialized niche that possibly supports such long-term cell fate plasticity.

## Main

In the embryo, stem cells and progenitor populations drive organogenesis by restricting their lineage potentials and acquiring specialized fates as development progresses to eventually give rise to all mature terminally differentiated cell types^9^. By contrast, tissue-resident adult stem cell populations contribute to tissue homeostasis and repair by replenishing specialized differentiated cells^10^. Recent single cell-based analyses have started to unveil a tremendous cellular heterogeneity in both embryonic progenitor and adult stem cell compartments^1–3, 11^. These studies suggest that progenitor populations with varying degrees of lineage commitment co-exist with cells harbouring long-term cell fate plasticity within the same embryonic tissue or stem cell compartment^3, 5, 12^. For instance, single cell-based studies have led to a revised road map for lineage commitment in the hematopoietic system, challenging the demarcations between stem cell and progenitor populations and the timing of cell fate choices in the hematopoietic lineage^3, 4^. Nevertheless, single cell-based data present static snapshots of cell states^13–15^, hampering sometimes to assess temporal and spatial relationships between concurrent progenitor populations, their progeny, and their functional relevance for organ development.

Here, we investigated lineage dynamics and relationships within the hepato- pancreato-biliary progenitor compartment, which represents a paradigm for organogenesis, whereby a small pool of progenitor gives rise to a variety of different adult tissues, including liver, pancreas, gallbladder, and extra-hepatic bile ducts^6, 7, 16^. Combining lineage tracing analyses with quantitative modelling approach and single-cell transcriptomics, we show that a long-term multipotent progenitor domain persists in the pancreato-biliary organ rudiment, contributing to the formation not only of the pancreas and gallbladder but also the liver. Moreover, we define a specialized niche that possibly supports such long-term cell fate plasticity at the border between liver and pancreas.

### Long-term plasticity between hepatic and pancreato-biliary cells

Hepato-pancreato-biliary organogenesis has been primarily described as a process of sequential binary cell fate decisions. First, a common endoderm progenitor population residing within the anterior ventral gut tube (*a.k.a.* ventral foregut) undergoes cell fate segregation into hepatic and pancreato-biliary progenitors around embryonic day (E) 8.5 in the mouse^6, 7, 17, 18^. Subsequently, the pancreato- biliary progenitors give rise to ventral pancreatic and gallbladder progenitors^6, 19, 20^. In addition, a separate endodermal progenitor population, which arises from the dorsal side of the anterior gut tube, also contributes to the pancreas^7, 21^. To quantitatively assess the tissue dynamics accompanying the ontogeny of the hepato-pancreato- biliary organ system, we performed immunofluorescence (IF) staining for markers of liver (Prox1)^22^ and pancreato-biliary (Prox1, Pdx1, Sox17)^20, 21^ cell types in mouse embryos between E8.5 [equivalent to 10 somite stage (ss)] and E11.5 (45 ss) and measured the number of cells and volume of each organ rudiment (Fig. 1a). The measurements documented a considerably faster increase in size of the liver bud as compared to the pancreato-biliary bud (Fig. 1b, Extended Data Fig. 1a,b). To determine whether the profound difference between liver and pancreato-biliary buds is due to different proliferative activity, we evaluated the mitotic marker phospho-histone 3 (pH3) in the different cell populations. Unexpectedly, pH3 levels were higher in pancreato-biliary rudiments as compared to liver buds at early stages (E8.5), while no significant differences between the two organ rudiments were observed at later stages (Fig. 1c, Extended Data Fig. 1c,d). Consistently, *in vivo* BrdU incorporation assays showed comparable proliferative activity in hepatic and pancreato-biliary progenitor cells (Fig. 1d, Extended Data Fig. 1g,h). Moreover, no signs of apoptosis were detected in the pancreato-biliary rudiments at these embryonic stages^23^ (Extended Data Fig. 1i). Taken together, these results suggested that the differences in size between hepatic and pancreato-biliary organs are not due to higher levels of proliferation in the liver or increased apoptosis in the pancreato- biliary bud.

**Figure 1.**
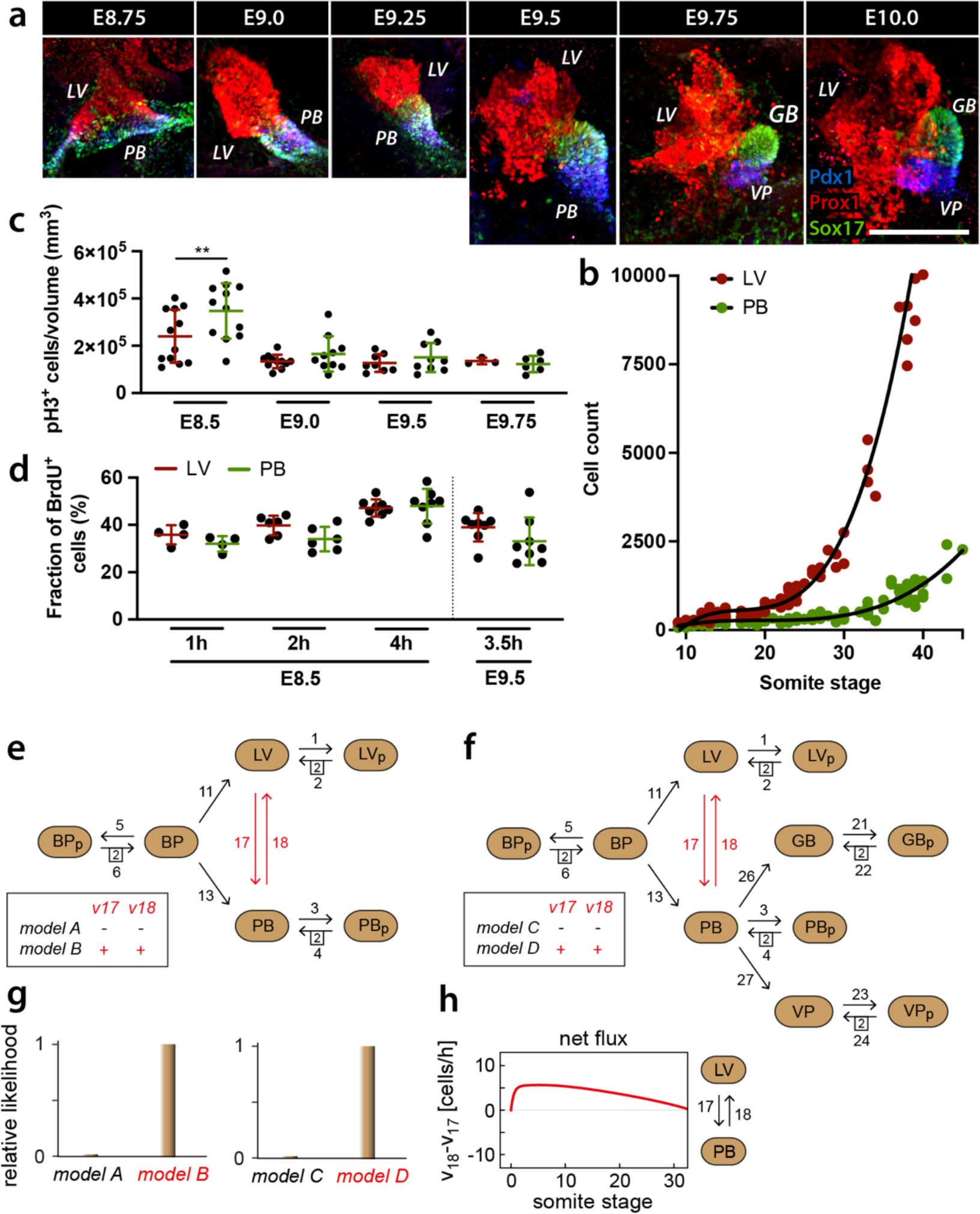
Modelling of tissue dynamics in hepato-pancreatic organ rudiments indicates long-term plasticity between hepatic and pancreatic cell fate. (a) Representative whole-mount immunofluorescence (IF) staining of embryos between E8.75 and E10.0 [12–28 somite stages (ss)] for the indicated markers. Prox1 (red) marks liver (LV) progenitors; Prox1, Pdx1 (blue), and Sox17 (green) mark pancreato-biliary progenitor cells (PB). After E9.75 (26 ss), the PB bud separates into two distinct organ domains, including the gallbladder (GB; Prox1/Sox17-double positive) and ventral pancreatic rudiments (VP; Prox1/Pdx1-double positive). Scale bar, 100µm. (b) Scatter plot showing cell counts in LV and PB buds at indicated somite stages. The number of progenitor cells was quantified on IF of cryosections of LV and PB buds stained with antibodies against Prox1, Pdx1 and Sox17. LV bud, n=72 embryos, PB bud, n=95 embryos. (c) Quantification of proliferation on whole-mount IF of LV (n=34 embryos) and PB buds (n=38 embryos). The number of pH3^+^ cells was normalized to the organ volume (mm^3^). Error bars represent ± s.d. One-way ANOVA test, p-value=0.006. (d) BrdU uptake in LV and PB progenitors of E8.5 and E9.5 embryos following indicated BrdU pulse-chasing periods (Extended Data Fig. 1g). Dot graphs summarize the fraction of BrdU-labelled cells relative to the total number in each respective organ rudiment (% of total cell count) examined at E8.5 and E9.5. Each dot represents the mean from an individual embryo [E8.5: n(1h)=4, n(2h)=6, n(4h)=8; E9.5: n(2h)=2, n(4h)=6]. Error bars represent ± s.d. One-way ANOVA test, ns. (e, f) Schematic representation of models A, B (e) and models C, D (f). Populations considered in these models include bipotent endoderm (*a.k.a.* ventral foregut) progenitor cells (BP), hepatic (LV), pancreato-biliary (PB), gallbladder (GB), and ventral pancreatic (VP) progenitors (Supplementary Information Note 1). (g) Comparison of relative likelihood, as determined by Akaike weights, of the different models identifies model B and D as better fitting the experimental data than models A and C, respectively. (h) Analysis of the temporal dynamics of flux differences (v_18_- v_17_) between LV and PB populations in model D shows a higher flux of PB cells to LV until about 30ss.

We then asked whether dynamic heterogeneity within the common progenitor pool may explain the differences between organs, whereby asynchronous fate-restriction or long-term cell plasticity might contribute to the rapid expansion of the liver while attenuating pancreato-biliary growth. To test this hypothesis, we employed a computational modelling approach that described the temporal changes of the sizes of the different progenitor populations (Supplementary Information Note 1).

We developed two sets of computational models, which differ in their complexity (Fig. 1e,f). The first set of models (models A and B) described the changes in the population size of three cell types, namely the bipotent endoderm progenitor (BP) cells, hepatic (LV) and pancreato-biliary (PB) progenitor cells (Fig. 1e). Each cell type is represented by two sub-populations: one sub-population of cells undergoing proliferation (pH3^+^), referred to as BPp, LVp, PBp, and one sub-population of cells that may differentiate or undergo the next proliferation cycle, referred to as BP, LV, PB. We incorporated two additional processes (Fig. 1e; red arrows 17 and 18) into the structure of model B that are missing in model A. These processes allow for cell fate plasticity between LV and PB progenitors, implying potential differentiation of both cell types into one another. To estimate the model parameters, model A and model B were fitted independently to the same experimental datasets. The experimental datasets included the total number of cells and pH3-positive (^+^) proliferating cells in LV and PB organ rudiments of embryos between E7.5 and E11.5, equivalent to 0-45 ss (Figs. 1b,c, Extended Data Fig. 2; Supplementary Table S1). We identified the model that best represented the experimental datasets by calculating the relative likelihood of models A and B^24^. The comparison of relative likelihood showed that the model allowing for cell fate plasticity (model B) explains the experimental data much better than the model without cell fate plasticity (model A) (Fig. 1g, Extended Data Fig. 2a).

**Figure 2.**
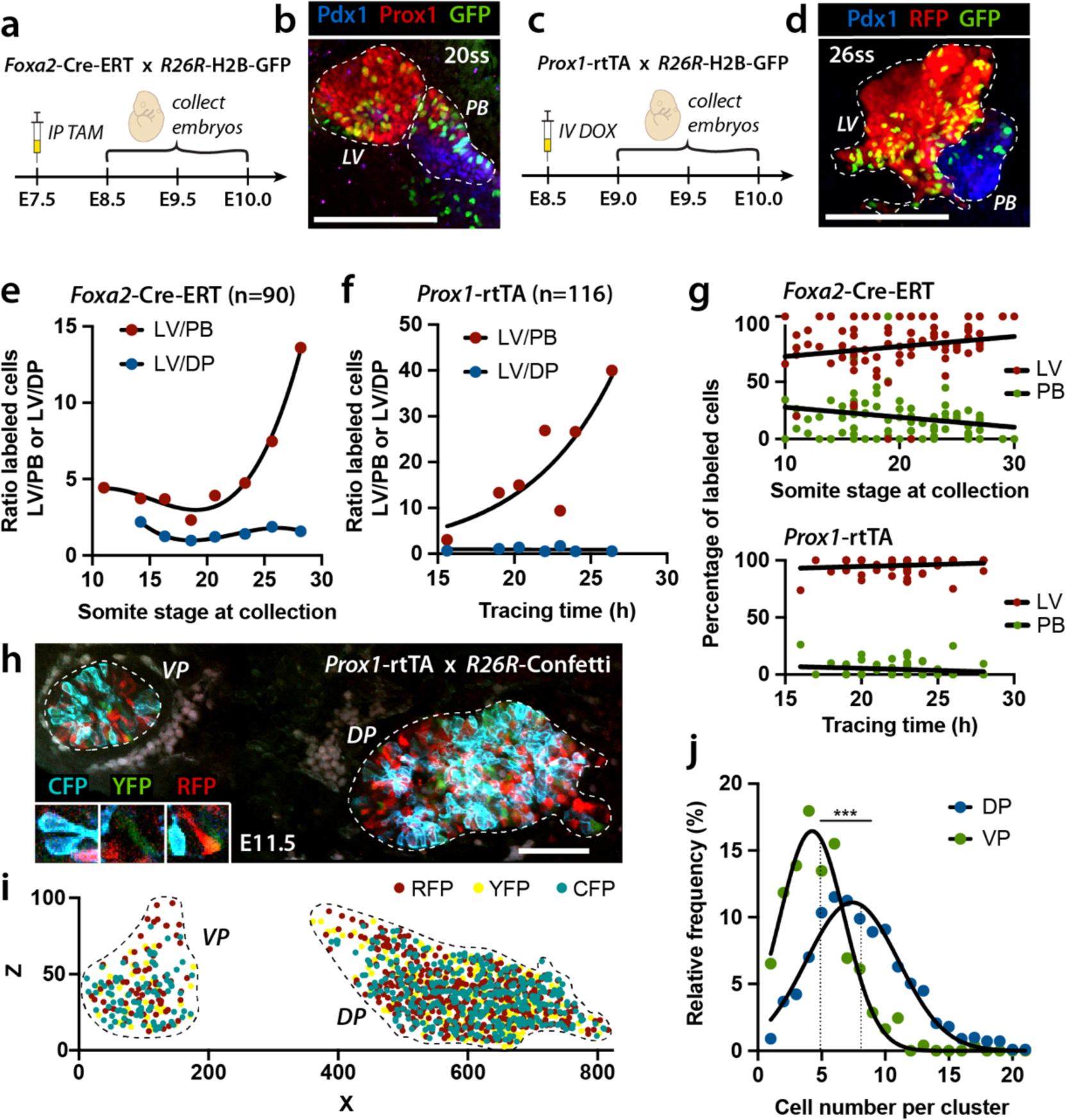
Lineage tracing of hepato-pancreato-biliary progenitor populations. **(a)** Schematic representation of breeding strategy and experimental setup to obtain Tg(*Foxa2*-Cre-ERT; *R26R*-H2B-GFP) embryos. Pregnant females were intraperitoneally (IP) injected with a single dose of tamoxifen (TAM; 12µg/g body weight) at E7.5. Labelled embryos were collected between E8.5-E10.0. **(b)** Representative whole-mount IF image of Tg(*Foxa2*-Cre-ERT;*R26R*-H2B-GFP) embryos at the indicated somite stage. Prox1 (red) marks liver (LV) progenitors, Prox1/Pdx1 (blue) double-staining marks pancreato-biliary (PB) progenitor cells. Dashed white lines demarcate LV and PB buds. GFP (green)-labelled cells were present in both LV and PB buds, even though a reduction in density of GFP^+^ cells was observed in the PB with increasing somite stage [around 29 ss; see (e)]. Scale bar, 100µm. **(c)** Schematic representation of breeding strategy and experimental setup to obtain Tg(*Prox1*-rtTA; *R26R*-H2B-GFP) embryos. Pregnant females were intravenously (IV) injected with doxycycline (DOX; 150µg/g body weight) at E8.5. Labelled embryos were collected between E9.0-E10.0. **(d)** Representative whole- mount IF image of E9.5 Tg(*Prox1*-rtTA; *R26R*-H2B-GFP) embryos. GFP (green)- labelled cells were present in both LV and PB buds, even though a reduction in density of GFP^+^ cells was observed in the PB with increasing tracing period [about 28h; see (f)]. Scale bar, 100µm. **(e, f)** Scatter plots showing the ratio of genetically labelled cells in LV to PB (LV/PB) or LV to dorsal pancreas (DP) against somite stage at the time of collection (e) or the length of the tracing period (f). Embryos of similar somite stage at time of collection (e) or tracing period (f) were grouped [(e) n=90 embryos; (f) n=116 embryos]. In (e), the shift in LV/PB ratio against somite stage is statistically significant (linear regression t-*test*, p-value=0.048). In (f), the shift in LV/PB ratio against tracing period is statistically significant (linear regression t-*test*, p-value=0.019). The ratio of labelled LV to labelled DP cells is not influenced by somite stage [(e) p=0.99] or tracing period [(f), p=0.79]. **(g)** Plots displaying the number of genetically labelled LV or PB progenitor cells, as percentage (%) of total labelled cells in both rudiments, against somite stage in *Foxa2*-Cre-ERT experiments (upper panel) or tracing period in *Prox1*-rtTA experiments (lower panel). Linear regression t-*test*, p-value=0.006 measured for *Foxa2*-Cre-ERT lineage tracing dataset; linear regression t-*test*, p-value=0.061 for *Prox1*-rtTA lineage tracing dataset. **(h)** Representative optical section of two-photon microscopy 3D scans of E11.5 Tg(*Prox1*-rtTA; *R26R*-Confetti) embryos. Genetically labelled cells expressing CFP, YFP, RFP, were detected in both ventral (VP) and dorsal pancreatic (DP) buds (circled by dashed white lines). Scale bar, 100µm. **(i)** *In silico* re-construction of genetically labelled cells in pancreatic buds of Tg(*Prox1*-rtTA;*R26R*-Confetti) embryos. Spot detection analysis in Imaris software was used to identify *xyz*- coordinates of individual genetically labelled cells and reconstruct the labelled tissues. **(j)** Clone size distribution established from Confetti lineage tracing experiments. Information on *xyz*-coordinates of labelled cells in DP and VP (i) was used for clustering individual cells into clonal clusters based on their geometric distance from one another^29^. Significantly different distribution of clone sizes was measured between VP and DP, with lower number of cells per clone in VP compared to DP. The vertical dotted lines indicate mean values; Mann-Whitney test, p-value <0.001.

Next, we increased the level of detail in the models to better predict the behaviour of individual hepato-pancreato-biliary sub-populations during organogenesis. To this aim, we developed a second set of computational models (models C and D) that extended the models A and B by including gallbladder (GB) progenitors and ventral pancreatic (VP) progenitors (Fig. 1f). Specifically, models C and D describe a population of BP progenitor cells that might differentiate into either LV (process 11) or PB progenitor cells (process 13). PB cells in turn might further differentiate into GB (process 26) or VP progenitors (process 27) (Fig. 1f). In addition, model D allows for cell fate plasticity between LV and PB progenitor populations through processes 17 and 18 (Fig. 1f). As in models A and B, the new models C and D take into account that each progenitor population has a sub-population of pH3^+^ proliferating cells, referred to as BPp, LVp, PBp, GBp, VPp. The GB progenitors were defined as Sox17^+^ cells^20^, and VP progenitors were defined as Pdx1^+^ cells^7, 20, 21^ (Supplementary Information Table S1). Specifically, to collect quantitative data on the separation of the PB progenitors into Sox17^+^ or Pdx1^+^ cells, we measured nuclear fluorescence intensity (FI) in individual cells in embryos stained for Pdx1 and Sox17 between E8.5 and E11.5 (Fig. 1a, Extended Data Fig. 3). Based on relative expression levels, cells were categorized as GB progenitors (Sox17^high^), VP progenitors (Pdx1^high^) or PB progenitors co-expressing Pdx1 and Sox17. The switch between the two states (Pdx1/Sox17-double positive or -single positive) occurred around E9.75 (Extended Data Figs. 3a,g-i,k).

**Figure 3.**
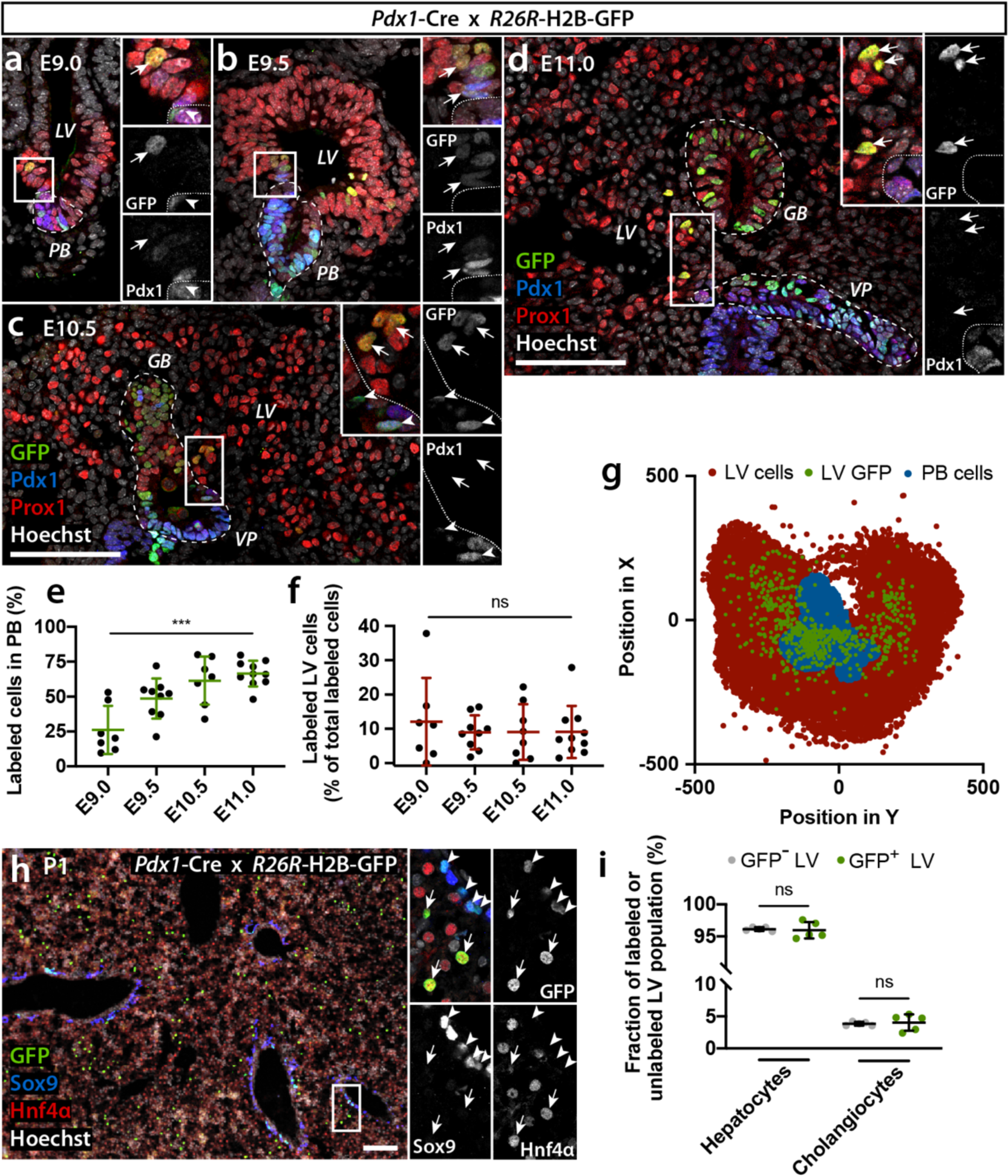
Pdx1-descendant cells are found in the liver. **(a-d)** Representative IF images of Tg(*Pdx1*-Cre; *R26R*-H2B-GFP) embryos at E9.0 (a), E9.5 (b), E10.5 (c), and E11.0 (d). GFP (green) identifies labelled cells descended from Pdx1^+^ progenitors; Prox1 (red) marks liver (LV), Prox1 and Pdx1 (blue) the pancreato- biliary organ buds (PB, demarcated by a white dashed line). Insets show higher magnification of boxed regions as merged and single GFP and Pdx1 channels. Arrows indicate GFP^+^ LV cells at the border with the PB (b) or with ventral pancreas (VP) and gallbladder (GB) (c,d); arrowheads indicate GFP^+^/Pdx1^+^ progenitors in PB (a) and VP (c). Tissues are counterstained with Hoechst dye. Scale bars, 100µm. **(e)** Quantification of the GFP^+^ PB progenitor population as percentage (%) of the total PB population in genetically labelled embryos (E9.0, n=7; E9.5, n=9; E10.5, n=8; E11.0, n=10) shows an increase at later embryonic stages (ANOVA followed by Kruskal-Wallis test, p <0.001). **(f)** Quantification of the GFP^+^ LV population as percentage (%) of the total labelled cell population in the ventral foregut at indicated time points (E9.0, n=7; E9.5, n=9; E10.5, n=8; E11.0, n=10). No statistically significant differences are detected among different embryonic stages (ANOVA followed by Kruskal-Wallis test, p =0.98). **(g)** Spatial representation of *Pdx1*-Cre lineage tracing experiments. IF of cryosections from E11.0 Tg(*Pdx1*-Cre; *R26R*-H2B- GFP) embryos (n=10) stained for Prox1, Pdx1, and GFP were digitalized and the *xyz*-coordinates for all individual GFP^−^ (LV) and GFP^+^ liver cells (LV GFP), and pancreato-biliary cells (PB) were obtained. The plot shows labelled descendants of Pdx1^+^ cells (LV-GFP, green dots) throughout the LV bud (red). Notably, GFP^+^ LV cells are found at higher density in close proximity to the PB bud (blue). **(h)** Representative IF image of a cryosection from a newborn Tg(*Pdx1*-Cre; *R26R*-H2B-GFP) mouse liver showing Hnf4α (red), Sox9 (blue), GFP (green) and nuclear counterstaining with Hoechst (grey). Insets show higher magnification of boxed region as merged and single grey scale fluorophore channels. GFP^+^ cells are positive for hepatocyte (arrows) or cholangiocyte (arrowheads) markers. Scale bar, 100µm. **(i)** Quantification of GFP^+^ and GFP^−^ hepatocytes and cholangiocytes on sections of the newborn LV. Hepatoblasts descended from genetically labelled ventral pancreatic progenitors differentiate into hepatocytes and cholangiocytes at the same ratio as non-labelled hepatoblasts (ANOVA followed by Kruskal-Wallis test, Dunn’s multiple comparisons test, p >0.99).

Comparison of the relative likelihoods showed that model D reproduces the observed data better than model C (Fig. 1g). This result along with our similar observation for model A and B indicate that a model accounting for fate plasticity between LV and PB cells is superior to a model neglecting fate plasticity in explaining the experimental data (Fig. 1g). Importantly, while plasticity from LV towards PB fate was allowed in D, simulation of model D showed a positive net flux of cells from the PB progenitor population towards the LV population until E10 (Fig. 1h, Extended Data Fig. 2c, d). In conclusion, our quantitative modelling approach supports the hypothesis that PB progenitors retain long-term cell plasticity and may acquire a hepatic fate, contributing to the rapid expansion of the liver bud at early developmental stages.

### Lineage tracing of hepatic and pancreato-biliary cell progenitors

To experimentally validate the predictions of the computational modelling approach, we used genetic lineage tracing to label foregut endoderm cells prior to (E7.5) or during the initial fate segregation (E8.5) into hepatic and pancreato-biliary progenitors in the mouse. First, we used the tamoxifen-inducible transgenic (Tg) *Foxa2*-CreERT line^25^ (hereafter referred to as Foxa2-CreERT) in combination with the Tg(*R26R*-H2B-GFP)^26^ reporter line (hereafter referred to as H2B-GFP) to induce genetic labelling of endoderm cells starting from E7.5. Next, we performed whole-mount IF staining to visualize and quantify GFP^+^ labelled cells in LV and PB buds of Tg embryos at different time points between E8.5 and E10 (Fig. 2a, b, Extended Data Fig. 4b). We found that, as development progresses, the relative number of GFP-labelled cells in PB rudiments decreases, while the relative number of labelled cells in LV buds increases (Fig. 2b, g), resulting in a shift in the ratio of labelled LV to PB progenitors (Fig. 2e).

**Figure 4.**
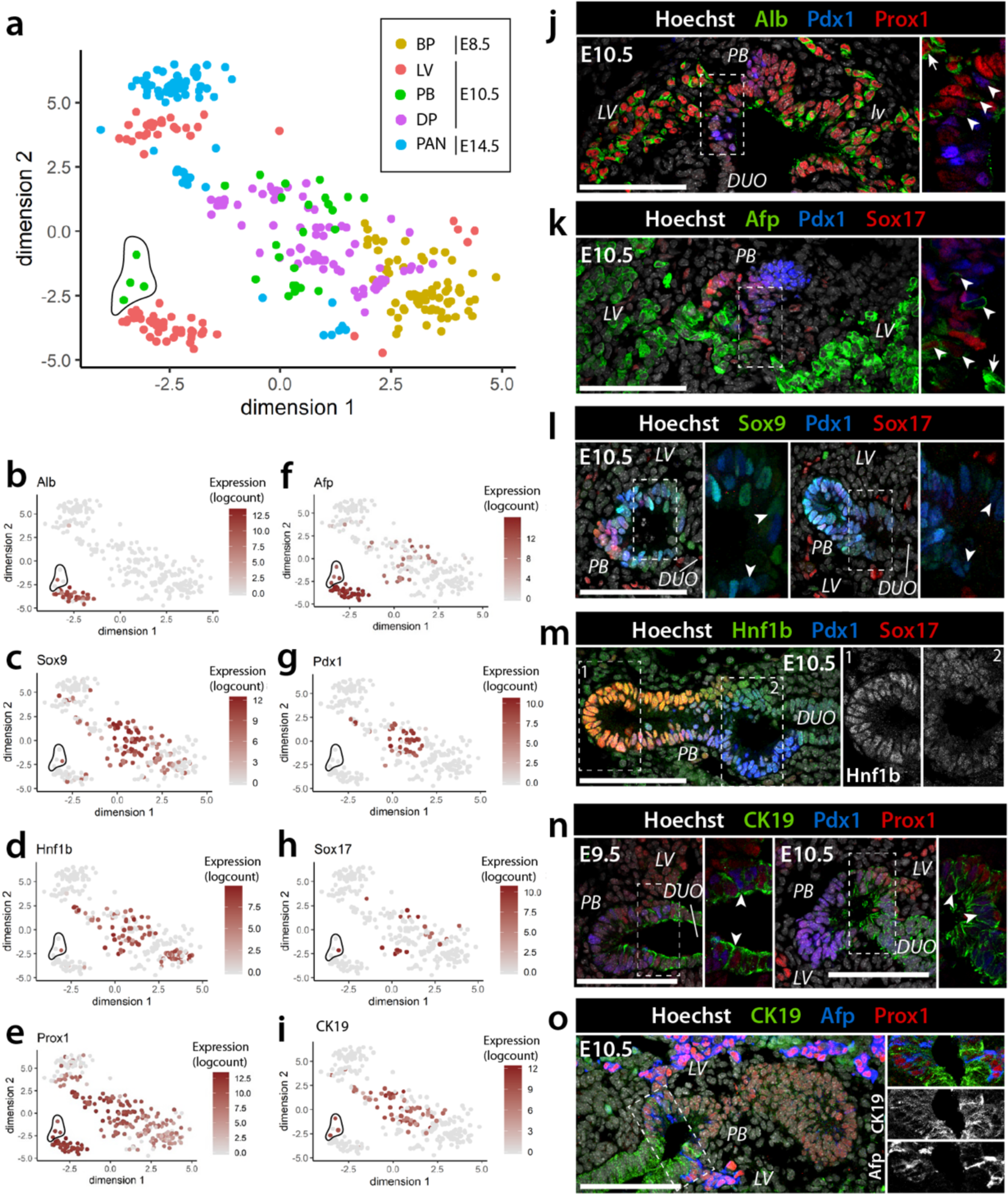
Sc-RNA-Seq identifies a distinct sub-population in the pancreato- biliary bud. **(a)** t-SNE plot visualization of sc-RNA-Seq dataset from E8.5 bipotent endoderm progenitors (BP, 82 cells), E10.5 hepatic (LV, 78 cells), dorsal pancreatic (DP, 86 cells), and ventral pancreato-biliary progenitor cells (PB, 27 cells), as well as E14.5 pancreatic cells (PAN, 73 cells). A subset of E10.5 PB cells clustered in close proximity to LV progenitors [circled by solid black line; referred to as intermediate progenitors (IMP)]. **(b-i)** Marker gene expression projected on t-SNE plots. Cells are projected into t-SNE space, as in (a), but are coloured by the relative expression of indicated hepato-pancreato-biliary marker genes instead of cluster assignment. Colours span a gradient from red (high expression) to grey (low expression). IMP cells are circled by a black line. **(j-o)** Representative IF images of E9.5 and E10.5 embryos stained for Prox1 (red; panels j, n, o), Pdx1 (blue; panels j-n) and Sox17 (red; panels k-m) and additional indicated markers. Right panels show higher magnifications of the boxed regions as merge (j-l, n-o) or single channels (m,o). In (j- k), arrowheads indicate cells at the border between LV and the PB (boxed region) positive for both hepatoblast markers [albumin (Alb) (j) or α-fetoprotein (Afp) (k)] and Pdx1. Of note, levels of hepatic markers in these cells are lower as compared to hepatoblasts (marked by arrows). In (l), low levels of Sox9 (green) and Pdx1 (blue) in cells (arrowheads) at the border between duodenum (DUO), LV, and PB buds (boxed region). In (m), Hnf1b is abundant in Sox17^+^ cells of the PB bud (boxed region 1) but low at the border between DUO, LV, and PB (boxed region 2). In (n,o) Cytokeratin 19 (CK19) displays high levels of expression in the border region (arrowheads) between DUO, LV, and PB buds (boxed regions) at E9.5 (left) and E10.5 (right). A subset of cells at the border are CK19^+^/AFP^+^ (o). Scale bars, 100µm.

Additionally, we performed genetic lineage tracing using an independent strategy based on a doxycycline-inducible Tg(*Prox1*-rtTA; *TetO*-Cre) mouse line (hereafter referred to as Prox1-rtTA) that we generated using BAC transgenesis (Extended Data Fig. 4; Supplementary Information Note 2). Compared to the Foxa2-CreERT, which is active in the entire endoderm, the Prox1-rtTA Tg strain enabled us to target the hepato-pancreatic endoderm^22, 27^ more specifically and at later time point, during the fate segregation between LV and PB progenitors (E8.5) (Fig. 2c, d; Extended Data Fig. 4a). In Prox1-rtTA;H2B-GFP Tg embryos we also observed a decrease in the relative number of labelled cells in the PB rudiment and a concurrent increase in the relative number of labelled LV cells with time (Fig. 2d, f, g), which is in line with the Foxa2-CreERT lineage tracing results (Fig. 2e). By contrast, the ratio between genetically labelled LV to labelled dorsal pancreatic progenitor cells, which arise from a separate progenitor domain in the dorsal endoderm, remained unchanged at different somite stage (Fig. 2e) or tracing time (Fig. 2f) in Foxa2-CreERT or Prox1- rtTA lineage tracing experiments, respectively. Thus, both lineage tracing results corroborated the quantitative analysis and computational simulations of the lineage, supporting a plastic relationship between LV and PB progenitors, with a potential gain of cells from the PB bud to the LV bud.

Next, we performed multicolour lineage tracing experiments using the Prox1-rtTA Tg line in combination with the Tg(*R26R*-Confetti)^28^ reporter line to define clonal dynamics in the developing pancreas. Genetic labelling was induced in mouse embryos at E8.5 and labelling events were assessed within both ventral (VP) and dorsal (DP) pancreatic progenitor pools at E11.5 (Fig. 2h, Extended Data Fig. 4), which showed comparable proliferation dynamics (Extended Data Fig. 1d-f). Specifically, we determined the number of labelled cells, their position in VP and DP organ buds and, then, performed clustering analysis to define the most probable separation of cells into clonal clusters^29^ (Fig. 2i, j). Notably, we found significant differences in clone size distribution between the two buds, with most clusters in the VP being composed of 4-5 cells, while about 8 cells per cluster were found in the DP (Fig. 2j). Given the similar proliferation rate between VP and DP progenitors (Extended Data Fig. 1d), the presence of smaller clones in VP is in line with the hypothesis that PB progenitors might separate and contribute to both LV and VP organ rudiments.

### Contribution of pancreato-biliary progenitors to the developing liver

To directly test the hypothesis of a contribution of PB progenitors to the LV bud, we performed additional lineage tracing experiments. Specifically, we used the Tg(*Pdx1*- Cre)^30^ line in combination with the H2B-GFP reporter line to induce genetic labelling in Pdx1^+^ progenitors in the embryo. IF staining detected GFP^+^ cells in the PB bud and both GB and pancreatic rudiments (Fig. 3a-d), as expected^20, 21^, but also documented GFP^+^ cells, descendant of Pdx1 progenitors, in the LV of Tg embryos (Fig. 3a-d). While the percentage of genetically labelled cell population in the pancreas and gallbladder increased with embryonic stage (Fig. 3e), the percentage of labelled cells found in the LV remained constant (average labelling index of 9.64% ± 1.4%) (Fig. 3f). Importantly, GFP^+^ cells were found in the LV throughout development as well as after birth with a constant fraction of labelled cells over total number of LV cells (Fig. 3, Extended Data Fig. 5d). *In silico* spatial reconstruction of confocal microscopy image stacks showed that genetically labelled cells are present throughout the LV bud, though the highest density was in close proximity to the PB rudiment at E10.5 and E11.0 (Fig. 3g, Extended Data Fig. 5a-c). Consistently, multicolour lineage tracing experiments using the Tg(Pdx1-Cre) line in combination with the Confetti reporter strain identified individual clones of the same colour distributed along hepatic chords (Extended Data Fig. 6).

**Figure 5.**
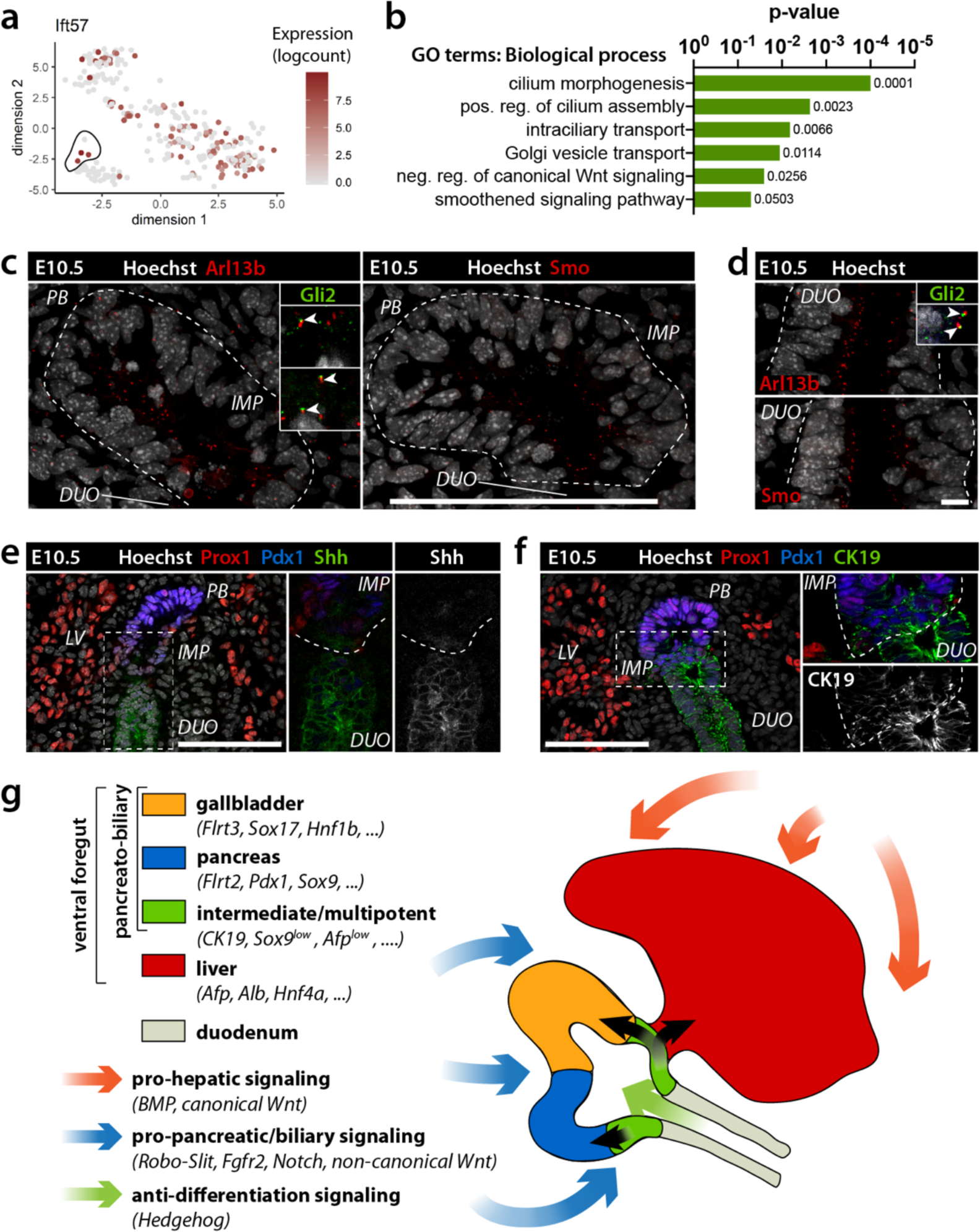
Unique signalling signature defines the intermediate progenitor domain bordering liver and pancreas. **(a)** tSNE plot of *Ift57* that was found to be enriched in E10.5 intermediate progenitors (IMP) (circled by a solid black line). **(b)** Biological process GO term analysis for genes enriched in E10.5 IMP cells compared to the hepatic (LV) and pancreato-biliary (PB) progenitor populations. **(c)** Representative IF images of E10.5 embryo cryosections stained for the primary cilia marker Arl13b (red) and Shh signalling pathway components Smoothened (Smo) (red) and Gli2 (green). Dotted white line demarcates the gut endoderm region at the border between LV, PB and duodenum (DUO), harbouring IMP cells. Arrowheads indicate localization of Gli2 at the tip of the cilia in a subset of IMP cells. Hoechst was used as nuclear counterstaining. Scale bar, 100µm. **(d)** Representative IF images of E10.5 duodenum stained for Arl13b, Gli2 and Smo. Arrowheads indicate localization of Gli2 at the cilia in the duodenum, a site of active Shh signalling^33^. The density of Smo^+^ primary cilia is highest in the duodenum and starts decreasing at the boundary with the PB, being still detected in cells at the border zone (c). Scale bar, 100µm. **(e, f)** Representative IF images of the E10.5 gut endoderm boundary region between LV, PB and duodenum (DUO) for the indicated markers. Shh (green) is abundant in DUO (e). CK19 staining (green) defines the border zone in (f), encompassing IMP cells. Right panels show higher magnifications of the boxed regions as merge or single channels of Shh (e) and CK19 (f) staining. Dotted line demarcates the border between IMP and DUO cells. Scale bars, 100µm. **(g)** Schematic representation of the hepato-pancreato-biliary region in E10.5 mouse embryo. The pancreato-biliary organ bud contains a progenitor domain (green) (IMP cells, green) that borders the liver rudiment and the duodenum with long-term plasticity, contributing to the pancreatic bud (blue), gallbladder (orange) or liver (red). Markers of the different progenitor domains and signalling pathways involved in hepato-pancreato-biliary development are listed on the left.

Finally, we further characterized GFP+ Pdx1-descendant cells in the LV and tested whether they express markers of both mature liver cell types (*e.g.* hepatocytes and cholangiocytes)^16^ after long-term lineage tracing experiments in Tg(Pdx1-Cre; H2B-GFP) transgenic newborn mice. Pdx1 expression was turned off in GFP^+^ cells in the LV (Fig. 3c,d) and the cells differentiated into both hepatocytes and cholangiocytes, being positive for Hnf4α and Glutamine Synthetase as well as Sox9 and Cytokeratin 19 (CK19), respectively (Fig. 3h, Extended Data Fig. 5e). Notably, quantification of the number of GFP^+^ hepatocytes and cholangiocytes showed that labelled progenitors contributed to the two cell types at the same ratio as non-labelled liver cells (Fig. 3i). Together these observations support a long-term contribution of Pdx1^+^ PB progenitors to the forming liver.

### Single-cell sequencing of the hepato-pancreato-biliary progenitor pool

Next, we asked whether long-term plasticity between LV and PB rudiments is confined to a specific progenitor pool. To answer this question, we performed single-cell RNA sequencing (sc-RNA-Seq) of E8.5 bipotent endoderm progenitors (BP; *a.k.a.* ventral foregut), E10.5 LV, ventral PB, DP progenitor cells and E14.5 pancreatic cells (PAN), isolated from Tg(Prox1-EGFP) embryos by FACS, as previously done for bulk-RNA-Seq^27^. After quality control (QC) and normalisation, we first applied dimensionality reduction to the dataset using a t-distributed stochastic neighbour embedding (t-SNE) clustering method to group cells by similarity in gene expression (Fig. 4a). As expected, E10.5 PB progenitors mostly clustered with E10.5 DP progenitor cells, lying between E8.5 BP and E14.5 PAN cells. Interestingly, a small subset of PB progenitor cells clustered in close proximity to E10.5 LV progenitors, possibly sharing a closer molecular signature with them (circled in Fig. 4a).

Importantly, the expression of well-known cell-type-specific markers perfectly correlated with the cluster allocation (Fig. 4b-i). For instance, hepatic marker genes (*e.g. Albumin*, α*-fetoprotein*) were strongly expressed in the LV cluster but not in the PB one, while pancreatic (*e.g. Pdx1, Sox9, Ptf1a*) and biliary (*e.g. Sox17*α, *Onecut1*, *Hnf1*β) markers were enriched in PB but not in LV cell clusters (Figs. 4b-h, Extended Data Fig. 7). By contrast, the sub-population of PB cells, which is close to the LV cluster, exhibited an intermediate molecular profile, displaying for example low expression of hepatic (*Afp*) and high levels of some pancreato-biliary markers (*CK19*) (Fig. 4, Extended Data Figs. 7, 8). IF staining for a selection of these markers validated the sc-RNA-Seq results and enabled us to define the spatial location of such intermediate progenitor pool (hereafter referred to as IMP) in the embryo.

Specifically, the combination of CK19 and AFP antibodies identified a small set of cells in the PB bud bordering liver and duodenum at E9.5 and E10.5 that are also positive for Prox1 (Fig. 4j, n, o) and express low levels of Pdx1 (Fig. 4j-n). The location of the IMP cells overlapped with the one of the genetically labelled cells after short-term lineage tracing experiments in Tg(Pdx1-Cre; H2B-GFP) transgenic embryos (Extended Data Fig. 5f), suggesting that this is the multipotent progenitor population contributing to both pancreato-biliary and hepatic lineages.

Next, we searched for signalling molecules that are enriched in the IMP pool and might be responsible for the long-term multipotency. Known pro-hepatic (*e.g.* BMP, canonical Wnt) and pro-pancreatic (*e.g.* Notch, Fgfr2, Robo-Slit, non-canonical Wnt) signalling pathway components were found enriched in LV and PB progenitor cells, respectively, as previously reported ^23, 27, 31, 32^ (Extended Data Fig. 9). We also identified several genes, which were not previously associated with LV and PB rudiments, such as regulators of primary cilium assembly and intraflagellar transport (*Ift57*, *Ift52*, *Ift43*), especially enriched in the small IMP population (Fig. 5a, b, Extended Data Figs. 7-9). Consistently, GO enrichment analysis revealed that the top enriched terms in IMP progenitors are related to primary cilium organization and regulation of Hedgehog signalling (*e.g.* “Cilium morphogenesis”, “Cilium assembly”, “Intraciliary transport”, “Smoothened signalling pathway”) (Fig. 5b).

The Hedgehog signalling is a well-known antagonist of pancreatic cell fate^7, 33^. We found high level of Shh ligand in the duodenum (Fig. 5e, Extended Data Fig. 8), which is in line with previous *in situ* hybridization^33^, and accumulation of Smoothened (Smo) in the primary cilia of the duodenum as well as IMP cells, but strongly decreased in PB progenitors (Fig. 5c, d). Increased ciliary Smo levels are associated with activation of Gli transcription factors^34, 35^. Co-localization of the primary cilia marker Arl13b with Gli2 in a subset of cilia tips in IMP cells at the border between the PB and LV buds (Fig. 5c) further corroborated active Hedgehog signalling in this location. Together these results suggest that the IMP population at the boundary the PB and LV buds receives Hedgehog signals from the neighbouring duodenum, which in turn might help to prevent differentiation and maintain long-term multipotency (Fig. 5g).

## Discussion

By combining experimental data with *in silico* modelling, we quantitatively characterized the hepato-pancreato-biliary tissue dynamics *in vivo* and identified a long-term progenitor population residing in the PB bud that contributes to pancreatic, biliary, and hepatic organ domains during development (Fig. 5). Thus, our findings indicate sustained cell fate plasticity between progenitor populations as an important feature underlying hepato-pancreato-biliary development, instead of occurring as sequential binary cell fate decisions, as previously suggested^6, 8, 18^.

Long-term cell fate plasticity between ventral foregut derivatives is supported by previous genetic studies in the mouse, in which ablation of transcriptional regulators or signalling effectors led to aberrant fate switches in the endoderm. For instance, deletion of *Sox9*^36^ or *Robo1*/*Robo2*^23^ genes induces hepatic fate in the ventral pancreatic organ domain. Similarly, deletion of *Sox17* results in the appearance of Pdx1^+^ cells in the liver bud^20^, while genetic deficiency for *Hes1* induces pancreatic fate in biliary tissues^37, 38^. In addition, fate switches have been reported in adult tissues upon injury, metabolic stress or in the context of rare cancer types^6, 8^, possibly recapitulating the developmental cell fate plasticity between hepatic and pancreato-biliary progenitors presented here.

Our findings suggest that the IMP pool resides in a specialized *niche*, which preserves them in an undifferentiated/multipotent state for longer time by concomitant low-level activation of mutually inhibitory pro-pancreatic, pro-hepatic, and Hedgehog signalling pathways. Hence, these progenitors retain the ability to undergo hepatic, pancreatic, or biliary differentiation upon leaving this specific environment. It is likely that tissue-specific multipotent progenitor *niche(s)* are not unique to the hepato-pancreato-biliary system but present in various developing organs. Maintaining multipotent cells in a developing organ may have a similar purpose as maintaining a stem cell *niche* in an adult tissue, providing concomitantly homeostatic regulation and improving resilience during organogenesis, for example following developmental delay or loss of a lineage-restricted cell population. Failure to maintain such a multipotent progenitor domain might result in human genetic syndromes, such as the Martínez-Frías syndrome^39^, Mitchell-Riley syndrome^40^, as well as congenital or environmentally caused human malformations, featuring multiorgan phenotypes in liver, pancreas and gallbladder^6, 8, 41^.

Finally, such long-term multipotent progenitor population might to some extent explain the rapid growth of the liver compared to the pancreato-biliary rudimentary despite their similar proliferation. A better understanding of such cell plasticity during development will shed light on the programs underlying the growth of these related organs. Moreover, further investigations are required to determine whether a subset of hepatocytes retains such plasticity also in adult life and could be targeted for improving stem cell programming or reprogramming strategies in regenerative medicine.

## Acknowledgments

We thank all the members of the Spagnoli laboratory for their useful comments and suggestions on the study. We thank Heike Naumann for technical help. We thank Gregory Pazour for IFT57 antibody (UMass, MA, USA) and Annabel Christ and Thomas Willnow for the Gli2 and Smoothened antibodies (MDC, Berlin, Germany). We thank the MDC Transgenic Unit for technical help in generating the Prox1-rtTA mouse strain. Maintenance of the two-photon microscopy setup was supported by the staff of the Advanced Light Microscopy technology platform via funding from the MDC in the Helmholtz Association. We thank Christian Beisel of the Genomics Facility of D-BSSE for NGS RNA sequencing. This research was supported by funds from the Helmholtz Association, European Union’s Horizon 2020 Research and Innovation Programme (Pan3DP Grant agreement no. 800981); D.W. was a recipient of a BIH (Tr. PhD) fellowship.

## Author contributions

F.M.S. and D.W. conceived the study, designed the experiments and wrote the manuscript. D.W. performed all the experiments. I.P. generated the Prox1-rtTA transgenic strain. A.M. helped with the Confetti lineage tracing experiments and two-photon image acquisition. U.B. and J.W. developed the mathematical modelling approach. M.L.V., A.V., Z.K. performed sc-RNA-Seq data analyses.

## Competing interests

Authors declare no competing interests.

**Correspondence and request for materials** should be addressed to FMS.

## Online Methods

### Mouse strains

Mice used in this study were on a C57BL/6 genetic background and kept under standard housing conditions. Transgenic mouse lines Tg(*Foxa2-* CreERT)^25^, Tg(*Pdx1*-Cre)^30^, Tg(*TetO*-Cre)^42^, Tg(*R26R*-Confetti)^28^, Tg(*R26R*-H2B- EGFP)^26^, and Tg(*Prox1*-EGFP)^27, 43^ were previously described. The Tg(*Prox1*-rtTA) transgenic mouse line was generated in the Spagnoli laboratory by modifying a mouse bacterial artificial chromosome (BAC) clone (RP23-360I16) spanning the *Prox1* gene and its upstream and downstream regulatory sequences. A transgenic expression cassette encoding for the reverse tetracycline-controlled transactivator (rtTA) fused to mCherry via a *2A* self-cleaving peptide sequence was inserted at the ATG of the *Prox1* open reading frame using homologous recombination^43, 44^. The transgene-modified BAC was subsequently introduced into the germline of C57BL/6 mice using pro-nuclear injection (see Supplementary Information Note 2). Embryos were staged according to Theiler staging criteria (E7.5) or by counting somites (E8.0- E11.5). For labelling of embryonic tissues, pregnant females were injected i.v. with doxycycline (Dox) or i.p. with tamoxifen (TAM) or BrdU using the indicated concentrations. All animal experimentation was performed after approval of protocols and according to the regulations of local authorities (Landesamt für Gesundheit und Soziales, Berlin).

### Immunohistochemistry

Embryos or newborn mice were dissected and fixed in 4% paraformaldehyde (PFA) in phosphate buffered saline (PBS) for 2h at room temperature (E7.5-9.5 embryos) or overnight at 4°C (E10.5-18.5 embryos, tissues). Fixed embryonic tissues were equilibrated overnight in 20% sucrose solution and embedded in O.C.T. compound (Tissue-Tek®, Sakura®). Cryosections (10 µm) were incubated with TSA (Perkin Elmer) blocking buffer for 1 hour at room temperature and afterwards with primary antibodies at the appropriate dilution (see Supplementary Table S6). If necessary, antigen retrieval was performed by boiling the slides for 20 minutes in citrate buffer (Dako). Hoechst 33342 counterstaining was used at a concentration of 250 ng/mL. Slides were mounted with Dako fluorescent mounting medium and imaged on a Zeiss LSM 700 confocal microscope using 40x, 63x oil or 10x water immersion objectives. For BrdU labelling, immunostaining with primary and secondary antibodies was followed by post-fixation with 4% PFA, treatment with 2.4M HCl for 30min at 37°C and immunodetection of BrdU.

For whole-mount IF staining, fixed embryos or embryonic tissue were blocked for at least 1h in PBSMT (2% milk powder, 0.5% Triton X100, in PBS). Samples were incubated overnight at 4°C with primary antibodies diluted in PBSMT (Supplementary Table S6), followed by overnight incubation at 4°C with secondary antibodies (1:500). Stained embryos were dehydrated in a methanol dilution series and clarified prior to imaging with methyl salicylate. Clarified embryos were imaged on a paraffin-sealed glass depression slide with a Zeiss LSM 700 confocal microscope using a 10x water immersion objective.

### Imaging native reporter fluorescence

To image native fluorescence of reporter proteins, fixed tissues were clarified in Sca*l*eA2 (4M urea, 0.1% Triton X-100, 10% glycerol, in H_2_O) for at least 2 weeks and up to 3 months at 4°C. Alternatively, tissues were sequentially clarified in CUBIC1 (25% wt/wt Urea, 25% wt/wt N,N,N’,N’- tetrakis(2-hydroxypropyl) ethylenediamine, 15% wt/wt Triton, in H_2_O) and CUBIC2 (50% wt/wt sucrose, 25% wt/wt Urea, 10% wt/wt 2,20,20’-nitrilotriethanol, 0.1% Triton, in H_2_O). The clarified tissues were mounted in paraffin-sealed CoverWell^TM^ imaging chambers and imaged using a LaVision BioTec TriM Scope II two-photon microscope (Bielefeld, Germany). Simultaneous excitation of CFP, GFP/YFP and RFP was obtained by combining 3 pulsed laser beams. CFP was excited at 860 nm using part of the output of a Chameleon Ultra II Ti:Sapph laser (Coherent, Santa Clara, CA, USA). The other part of this output was used to pump an optical parametric oscillator OPO (APE, Berlin, Germany) to obtain the 1100 nm wavelength for the RFP excitation. The GFP/YFP excitation was done at 940 nm using the beam of a Mira 900 Ti:Sapph laser pumped by a Verdi 5W laser (Coherent, Santa Clara, CA, USA). Images were acquired using a dedicated multiphoton water immersion 25x/1.10 NA objective (Nikon, Tokyo, Japan) by tile scanning areas of 250x250µm (318x318 pixels) spaced at 2 µm in *z*. The emission of the fluorescent proteins was separated using dichroic mirrors (long pass 495 nm and 560 nm) and detected via 3 bandpass filters (475/50 nm for CFP, 525/40 nm for GFP/YFP and 624/40 nm for RFP) using GaAs(P) detectors (Hamamatsu, Hamamatsu City, Japan).

### Image analysis

The open-source ImageJ/Fiji software (https://fiji.sc/) was used to measure the tissue area, organ bud volume and cell counts on cryosections. Liver cell counts of embryos older than E10.0 and newborn mice were obtained by using the automatic spot detection function of the Imaris software (Bitplane/Andor) with manual editing. Quantification of the fluorescence intensity and area on confocal images was done in ImageJ/Fiji using the region-of-interest (ROI) tool. For single nuclear fluorescence intensity quantification, intensity values in cells of interest were measured and corrected by linear normalization within each embryo. Cell coordinates for the analysis of Pdx1-Cre Tg lineage tracing were obtained using the spot detection function of Imaris software (Bitplane). The two-photon tile scans of Confetti embryos were stitched in ImageJ/Fiji using the ‘Grid/Collection stitching’ plugin^45^. To overcome the spectral cross-talk between CFP, YFP and RFP (GFP could not be identified based on its nuclear localization), the spectral unmixing plugin implemented in ImageJ/Fiji by J. Walter (https://imagej.nih.gov/ij/plugins/spectral-unmixing.html) was used. Coordinates of labelled cells were obtained by using the automatic spot detection function in Imaris with manual editing^46^. Clustering analysis of putative clones from the Confetti lineage tracing was implemented in the Hierarchical Clustering on Principal Components package^47^ and validation of clustering results was done using the NbClust package^29^, both available in the open source R software (https://.r-project.org/).

### Computational modelling

Temporal changes of the sizes of the cell populations are described by sets of ordinary differential equations (ODEs). We used the D2D Toolbox^48^ for Matlab (MathWorks, R2017b, The MathWorks Inc) to estimate the parameters of the ODE models. Simulations were performed using Mathematica 11 (Wolfram Research Inc.). Supplementary Dataset 1 presents the data used for computational modelling, while Supplementary Table S1 summarizes the attribution of markers Prox1, Pdx1, Sox17, and pH3 to each cell population (see Supplementary Information Note 1). Additional explanation on model analysis and detailed description of all models are provided in Supplementary Information.

### Single-cell RNA sequencing

Pancreatic and hepatic tissues from Tg(*Prox1*-EGFP) embryos (E8.5, E10.5, E14.5) were dissected in DEPC-PBS 1x and digested in 0.25% Trypsin in EDTA for 3’-5’ at 37 C (E8.5 to E10.5) or Collagenase for 10’ at 37 C (E14.5). Tissue digestion was stopped by adding DMEM medium with 10% FBS, cells were spinned 3’ at 300 g at 4°C, resuspended in DEPC-PBS and filtered through FACS tubes with Cell Strainer Cap (BD 352235) for immediate FACS sorting (BD *FACS Aria* II or III). Single cells were sorted into 96-well plates in lysis buffer (0.2% Triton X-100, 2U/µL Rnasin in nuclease-free water). Library preparation and transcriptome sequencing were performed by the Genomic sequencing facility at D-BSSE (Basel), according to the Smart-seq2 protocol^49^ (Invitrogen). cDNA profiles were checked on the Fragment Analyzer (AATI) and their concentration determined using Quant-iT PicoGreen dsDNA Assay Kit. Libraries were pooled and sequenced SR75 on an Illumina NextSeq 500 system (75 cycles High Output v2.5 kit). Reads were uniquely aligned to mouse reference genome (GRCm38) using STAR (v.2.5.3.a), by keeping only the reads that contain high confidence collapsed splice junctions. The gene expression matrices were generated employing GeneCounts as quantification type in STAR. The gtf and fasta formatted annotations for mouse were retrieved from GENCODE project (released version v26, primary assembly). Bioinformatic analysis was carried out using R statistical software and software packages from the open-source Bioconductor project^50^. QC was computed using Scater package^51^ and set to retain all samples in which at least 2000 genes and 500000 counts were observed. A gene filtering step (transcript > 2 in at least 2 cells) was also applied. Scran package was used for data normalization^52^. For data visualization, t-SNE plots with a perplexity value of 20% of the total number of cells were used. GO term analysis was conducted using David Bioinformatics tool.

### Statistical analysis

Data representation, statistical analyses, and plotting of regression lines were performed using GraphPad Prism and xls. Unless stated otherwise, data are shown as mean ± standard deviation (SD) and *n* numbers refer to biologically independent replicates. Statistical significance (p<0.05) was determined as indicated in figure legends using unpaired two-tailed Student t-test, Mann-Whitney test. For comparison between more than two samples, we performed one-way ANOVA followed by Kruskal-Wallis test (for datasets where we cannot assume a normal distribution), Brown-Forsythe and Welch test (for normal distributed datasets with different SD), with Dunn’s multiple comparisons test or Dunnett’s multiple comparisons test.

### Data availability

Single-cell dataset are available in GEO. All data supporting the findings of this study are available from the corresponding author on reasonable request.

---

**Extended Data Fig. 1.**
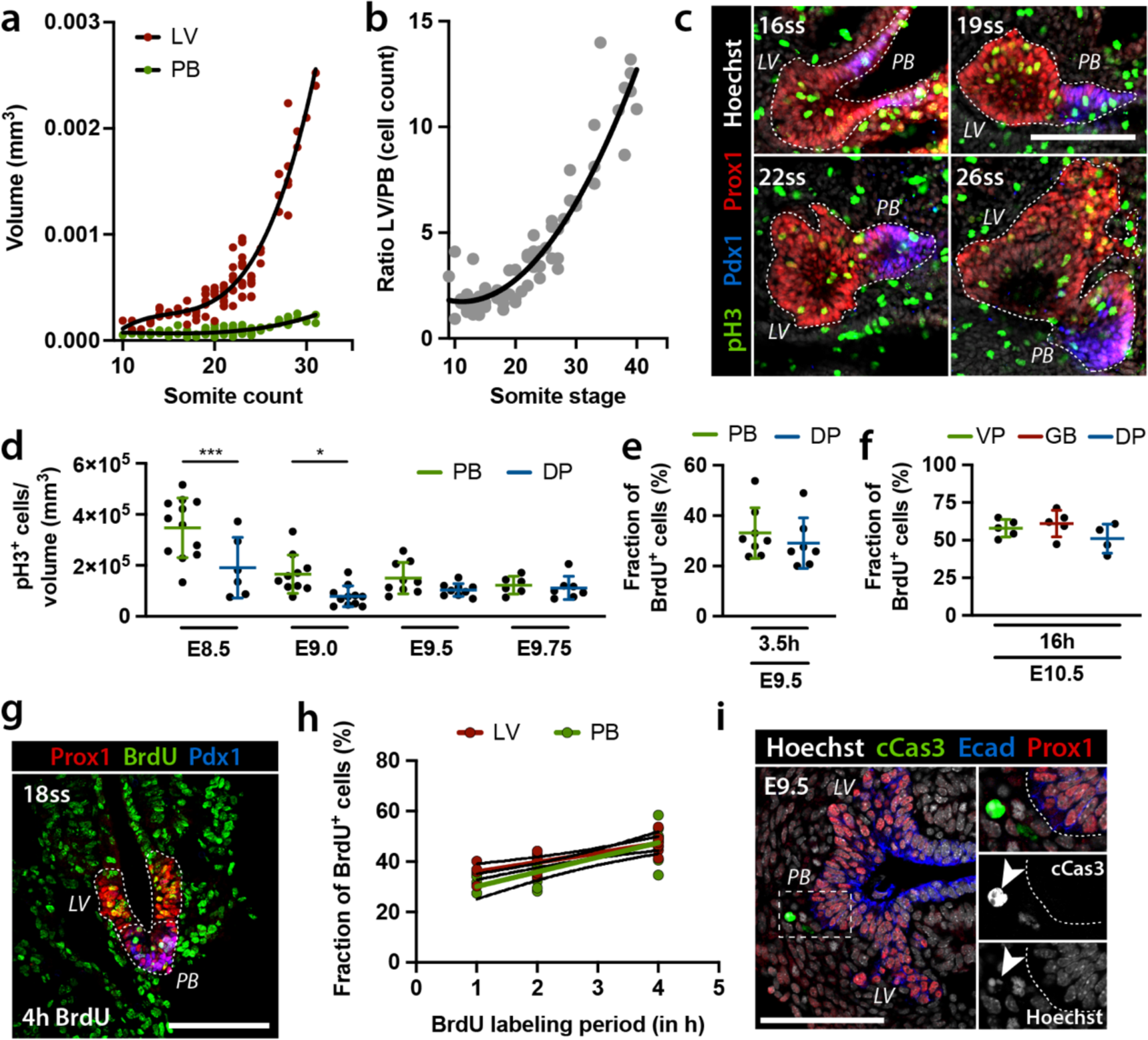
Proliferation dynamics in hepatic, ventral pancreato- biliary, and dorsal pancreatic organ rudiments. **(a)** Measurement of the liver (LV) bud (n=90 embryos) and pancreato-biliary (PB) bud (n=93 embryos) volumes at the indicated somite stages. 3D organ volume was reconstructed by measuring the surface area on individual optical sections of confocal z-series of whole-mount IFs (Fig. 1a); the average area measured in each embryo was then multiplied by tissue thickness. **(b)** Scatter plot showing the ratio of LV *versus* PB cell counts. Quantification of cell numbers in LV and PB buds was performed on IF stained cryosections (see Fig. 1b). **(c)** Representative whole-mount IF images of embryos at E9.0-E9.75 between 16-26 somite stages (ss) stained with antibodies against pH3 (green), Prox1 (red) and Pdx1 (blue). LV and PB buds are outlined by dashed white line. Scale bar, 100µm. **(d)** Quantification of pH3^+^ cells on whole-mount IF images of dorsal pancreas (DP) (n=33 embryos) and PB bud (n=38 embryos) normalized to organ volume (mm^3^). A statistically significant difference between the two progenitor populations is observed at E8.5 and E9.0, but not at later embryonic stages. Error bars represent ± s.d. One-way ANOVA test, p-value<0.001 (E8.5), p-value=0.04 (E9.0). **(e)** Fraction of BrdU^+^ cells in E9.5 embryos following the indicated pulse-chasing period is comparable in DP and ventral PB buds [n(2h)=2; n(PB, 4h)=6; n(DP, 4h)=5]. Two-tailed Mann-Whitney test, ns. **(f)** Fraction of BrdU^+^ cells in E10.5 embryos following the indicated pulse-chasing period [n=5 (GB); n=5 (VP); n=4 (DP)]. No statistically significant differences were detected between the different progenitor populations. One-way ANOVA test, ns. **(g)** Representative IF image of 18ss-embryo stained with the indicated antibodies showing BrdU labelled cells following a 4h-labelling period. Prox1 (red) marks LV progenitors, Prox1/Pdx1 (blue) double-staining the PB progenitors. LV and PB buds are outlined by a dashed white line. Scale bar, 100µm. **(h)** Graph illustrating the rate of cell proliferation as shown by the % of BrdU^+^ cells in E8.5 LV and PB organ domains at different labelling periods. An approximation of the cell cycle length (t_c_) was estimated based on BrdU incorporation using linear regression^53, 54^. LV slope = 3.8%/h; estimated cell cycle length = 26.6h (range with 95% confidence interval: 44.7h-19h). PB slope = 5.8%/h; estimated cell cycle length = 17.3h (range with 95% confidence interval: 29.6h- 12.2h). No statistically significant difference in average cell cycle length was measured between LV and PB progenitor cells (p-value=0.14). **(i)** Representative IF images of E9.5 embryo cryosections stained for Ecad, Prox1 and cleaved-Caspase3 (cCas3). No cCas3+ cells were found in the PB bud at these stages, while rare cells were detected in the surrounding mesenchyme and LV. Hoechst, nuclear counterstain. Insets show boxed area at higher magnification and cCas3 and Hoechst single channels. Arrowhead indicates apoptotic nucleus. Scale bar, 100rtm.

**Extended Data Fig. 2.**
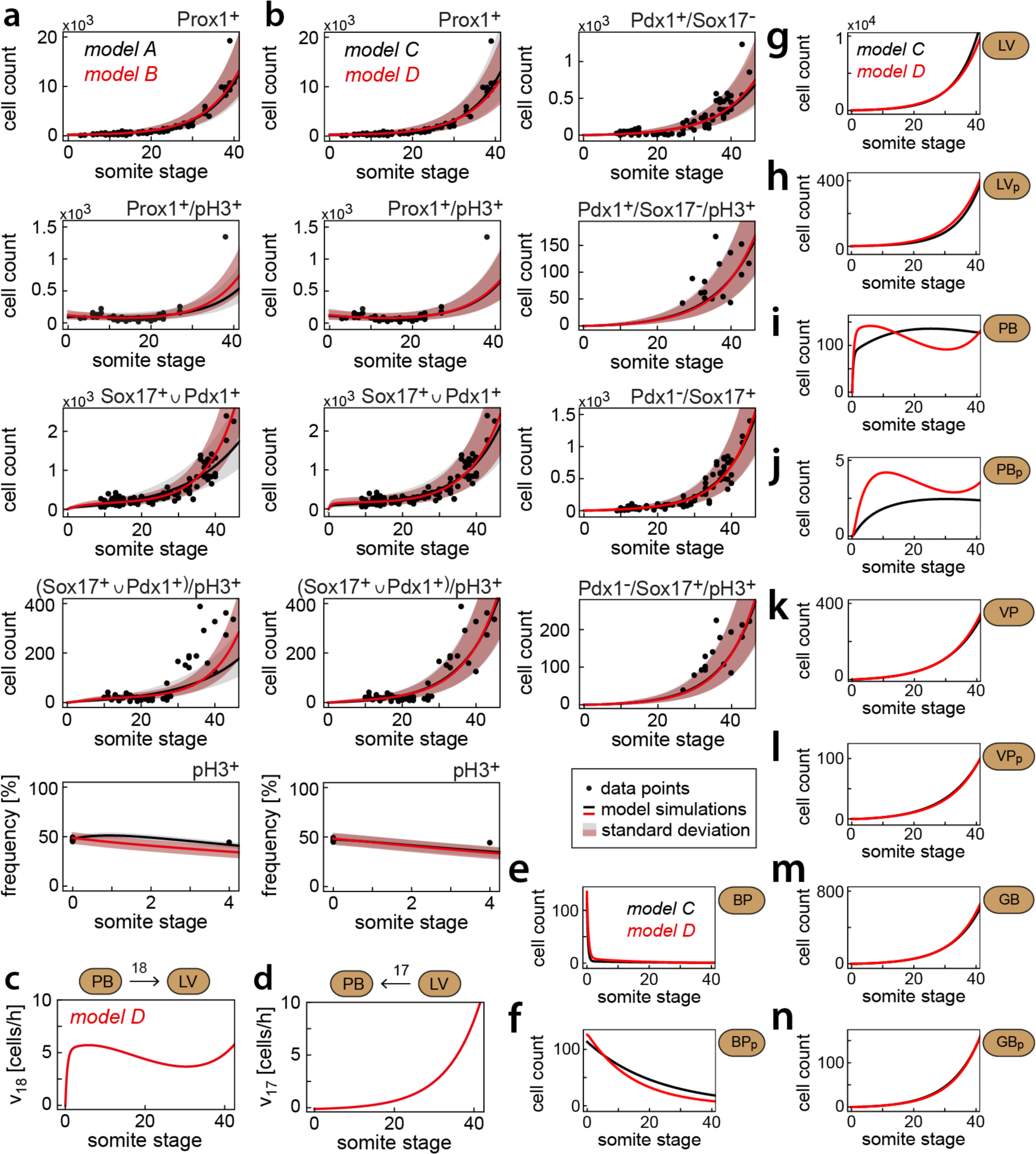
Extended computational modelling analysis of ventral anterior gut (*a.k.a*. foregut) endoderm development. **(a)** Fitting models A and B to experimental data. Best fit of models A (black solid line) and B (red solid line) and respective standard deviations are shown. Cell count data (black dots) of embryos collected between E7.5 and E11.5, equivalent to somite stages 0-45, was used to estimate the parameter values of the models. **(b)** Fitting models C and D to experimental cell count data (black dots). Best fit of model C (black solid line) and model D (red solid line) and respective standard deviations are shown. **(c, d)** Simulations of model D predict a flux of cells from the pancreato-biliary (PB) population to the hepatic (LV) one (c), and a delayed increasing flux in the reverse direction (d). **(e-n)** Simulations of the distinct cell populations (indicated above each panel) of model C (black solid line) and model D (red solid line).

**Extended Data Fig. 3.**
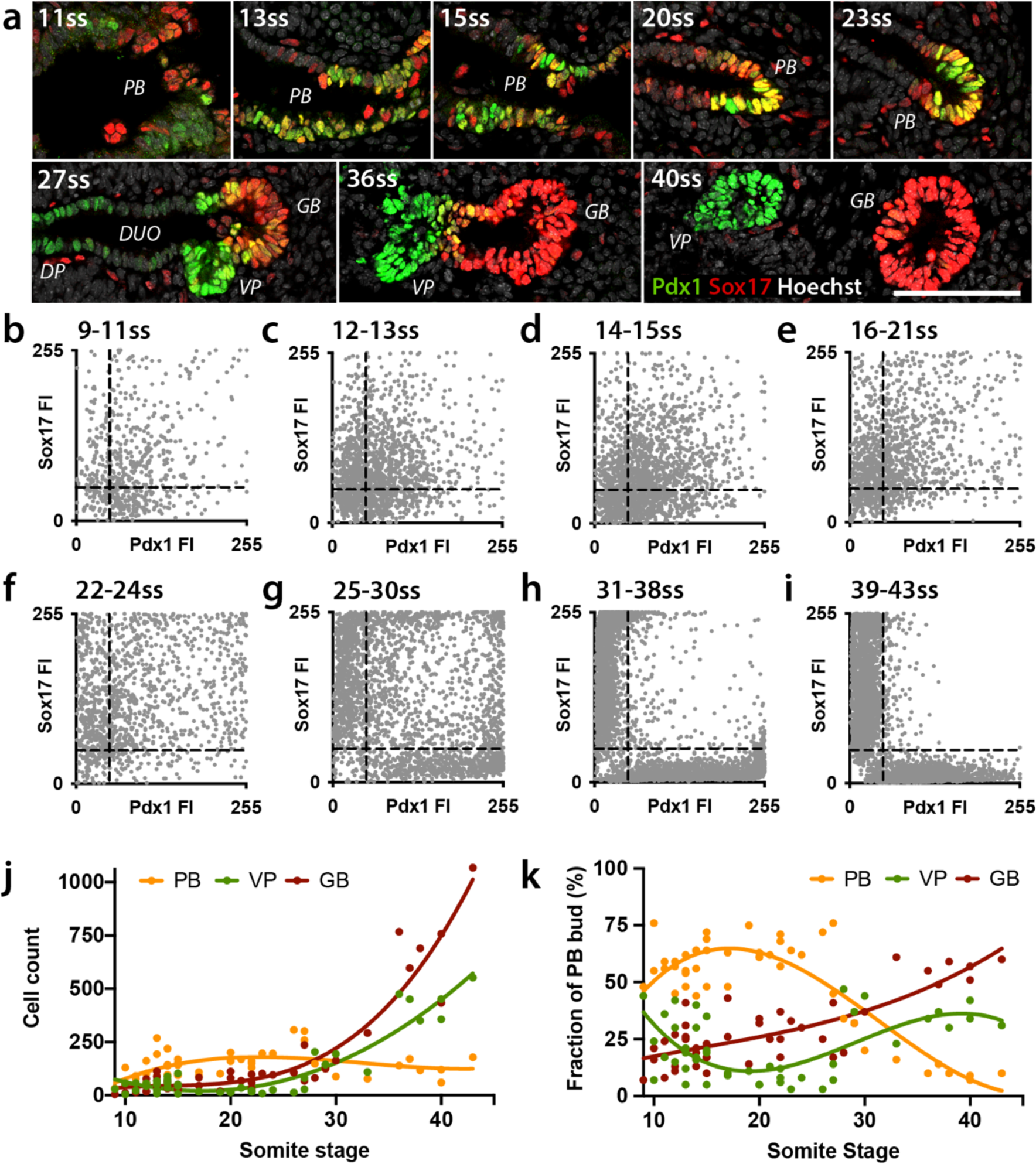
The pancreato-biliary bud contains cell subpopulations with distinct marker expression profiles. **(a)** Representative IF images of cryosections of the embryonic pancreato-biliary territory between E8.5 (11ss) and E11.0 (40ss). Pdx1 (green) and Sox17 (red) marks pancreato-biliary (PB) progenitor cells. After E9.75 (about 27ss), the PB bud segregates into gallbladder (GB; marked by Sox17) and ventral pancreatic bud (VP; marked by Pdx1). Hoechst was used as nuclear counterstain. Scale bar, 100µm. **(b-i)** Single-cell measurement of fluorescence intensity (FI) of Pdx1 and Sox17 in PB rudiments at E8.5-E11.0 (9- 43ss). Pdx1 and Sox17 FI values for individual cells were plotted. Embryos of similar somite stages were grouped as indicated (n, number of embryos; 9-11ss: n=5, 667 cells; 12-13ss: n=7, 1655 cells; 14-15ss: n=8, 1942 cells; 16-21ss: n=6, 1359 cells; 22-24ss: n=5, 1310 cells; 25-30ss: n=6, 2664 cells; 31-38ss: n=4, 4263 cells; 39-43ss: n=3, 3960 cells). From E8.5 to E9.5 (9ss-24ss), the majority of cells in the PB organ rudiment co-expressed Sox17 and Pdx1, while most cells at E10.0-E11.5 (31ss-43ss) acquired a Sox17^high^ or Pdx1^high^ identity. FI was measured with ImageJ/Fiji and values were corrected by linear normalisation within each embryo. Black dashed lines indicate sub-division of the progenitor populations based on Pdx1 and Sox17 FI levels. **(j, k)** Scatter plots displaying the number of cells in different PB rudiments against developmental time. FI data were used to categorize the subpopulations according to the relative expression levels of Pdx1 and Sox17. Pdx1^high^ sub-population corresponds to VP (Pdx1-FI>50, Sox17-FI≤50) (green); Sox17^high^ corresponds to GB (Pdx1-FI≤50, Sox17-FI>50) (red); Pdx1/ Sox17-double positive at low (Pdx1-FI≤50, Sox17-FI≤50) or high levels (Pdx1-FI>50, Sox17-FI >50) correspond to PB (yellow) progenitors. The three subpopulations exhibit distinct propagation kinetics as shown by plotting absolute cell counts (j) or fraction of subpopulation as percentage (%) of total cell number of the PB bud (k) against somite stage. While the relative fraction of PB progenitors decreased as compared to GB and VP progenitors after E9.75 (27ss), their total cell number remained constant throughout the analysed time period (j).

**Extended Data Fig. 4.**
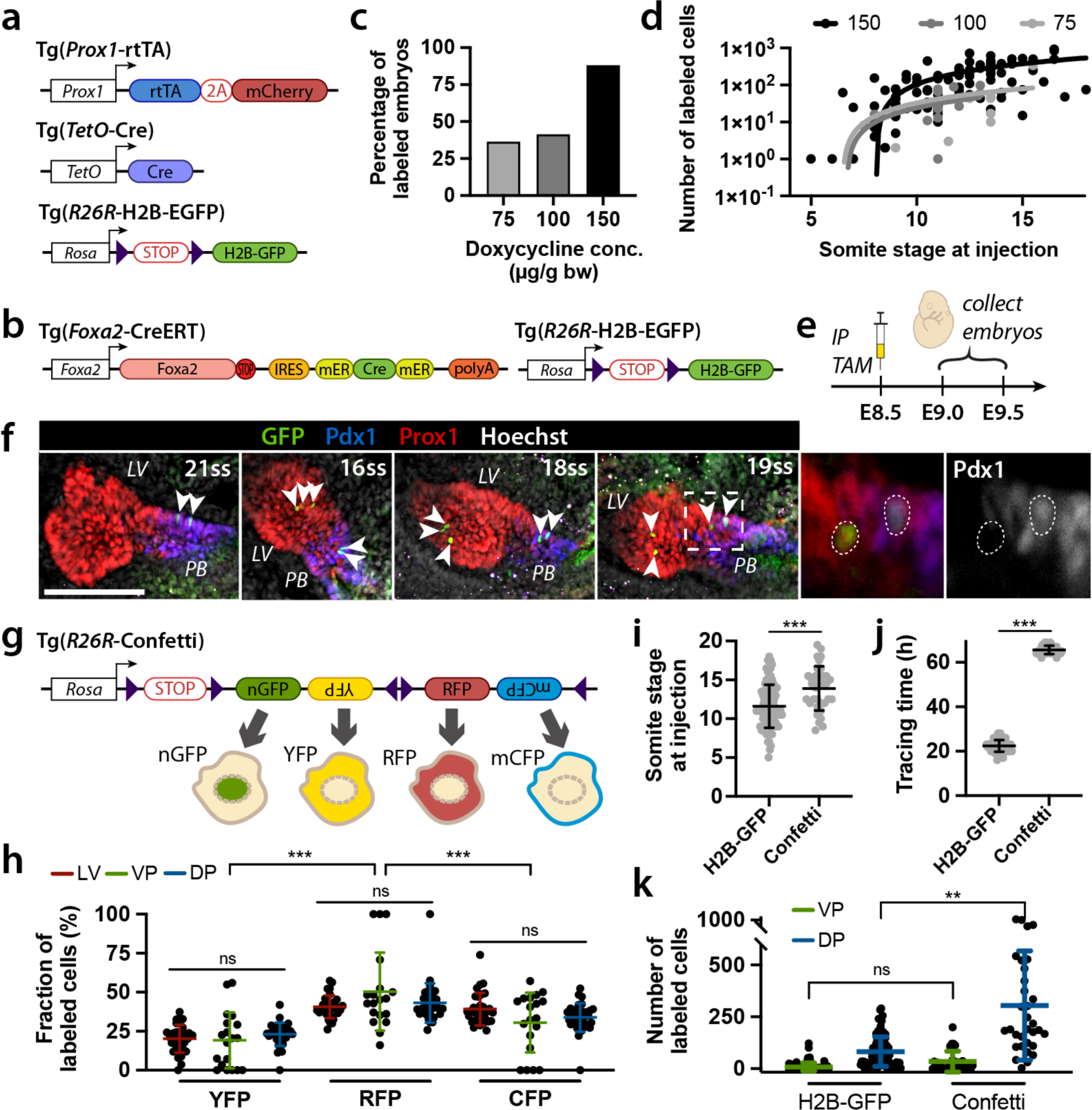
Extended lineage tracing analysis of hepato-pancreato- biliary progenitor populations. **(a, b)** Schematic representation of the transgenic constructs used for lineage tracing of hepato-pancreato-biliary progenitors. Abbreviations, IRES, internal ribosome entry site; mER, murine estrogen receptor. loxP sites shown as purple triangles. **(c, d)** Assessment of experimental parameters influencing Prox1-rtTA labelling efficiency. (c) Bar plot showing that % of embryos with induced label increase with increasing doxycycline doses (75µg/g bw, n=22; 100µg/g bw, n=29; 150µg/g bw, n=116). (d) Graph showing positive correlation between number of genetically labelled cells in individual embryos at the indicated doxycycline doses and time (somite stage) of labelling. For all further analyses, pregnant females were injected with 150µg/g bw of doxycycline. **(e)** Schematic representation of experimental setup for clonal labelling of Tg(*Foxa*-Cre-ERT; *R26R*- H2B-GFP) embryos. Pregnant females were intraperitoneally (IP) injected with a single dose of tamoxifen (TAM; 12µg/g body weight) at E8.5 and embryos collected at E9.0-E9.5 (n=15 embryos). **(f)** Representative whole-mount IF of Tg(*Foxa2*-Cre- ERT; *R26R*-H2B-GFP) embryos at E9.0-E9.5 (16-21ss) after TAM injection at E8.5. Arrowheads indicate GFP^+^ genetically labelled cells in liver (LV) and pancreato-biliary (PB) buds. In clonal labelling experiments, 2-3 labelled cells were often found in close proximity to one another, possibly being descendants of a common labelled progenitor. Boxed region is shown at a higher magnification in the right panels; white dotted circles indicate two adjacent cells, one Pdx1^+^ and the other one negative, which possibly arose from the same common progenitor. Scale bar, 100µm. **(g)** Schematic representation of the *R26R*-Confetti transgene. Following recombination, genetically labelled cells either express nuclear GFP (nGFP), cytoplasmic RFP or YFP, or membrane-associated CFP (mCFP). **(h)** Quantitative analysis of the contribution of the indicated fluorophores to the total number of labelled cells in the embryos shows characteristic induction efficiencies for each fluorophore that are comparable among different tissues but distinct among fluorophores [Welch ANOVA test, Dunnett’s multiple comparisons test, p <0.001]. **(i, j)** Assessment of the experimental parameters (i) somite stage at doxycycline injection and (j) tracing period in *Prox1*-rtTA lineage tracing experiments using the *R26R*-H2B-GFP (n=102 embryos) or the *R26R*-Confetti reporter (n=34 embryos) to genetically label embryos. Two-tailed Mann-Whitney test, p <0.001(i); p <0.001(j). **(k)** Quantification of total labelled cell numbers in dorsal (DP) and ventral (VP) pancreata of Tg(*Prox1*- rtTA; *R26R*-H2B-GFP) and Tg(*Prox1*-rtTA; *R26R*-Confetti) embryos. The number of labelled cells in DP but not in VP increased significantly with prolonged tracing period (mean tracing time: 22.4h for H2B-GFP, 65.6h for Confetti) [nonparametric Kruskal-Wallis test, followed by Dunn’s multiple comparisons test, p-value= 0.002 (DP), p-value= 0.08 (VP)].

**Extended Data Fig. 5.**
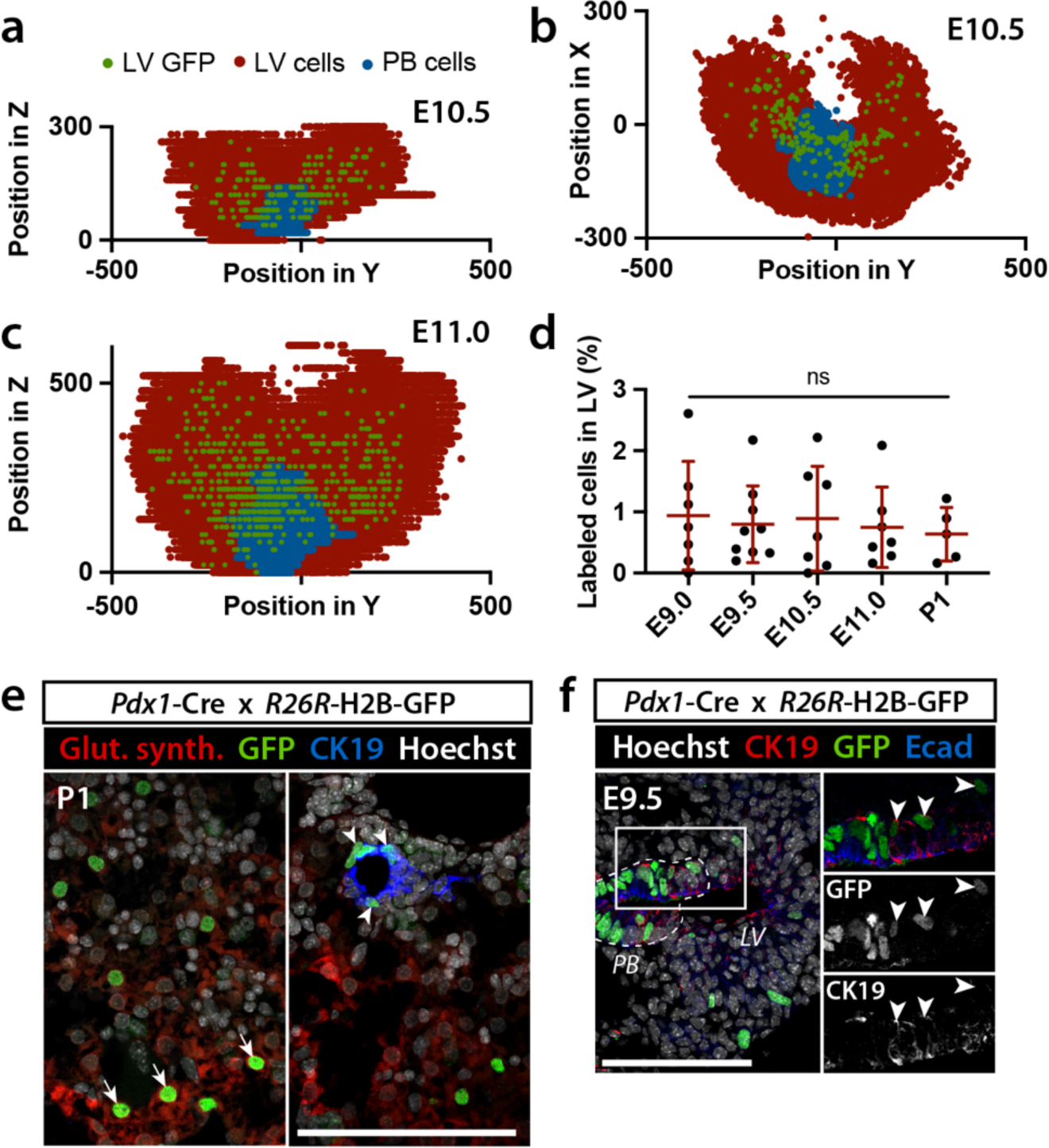
Analysis of lineage tracing of Pdx1^+^ pancreato-biliary progenitors. **(a-c)** Spatial representation of *Pdx1*-Cre lineage tracing experiments. IF images of cryosections from E10.5 (a, b) and E11.0 (c) Tg(*Pdx1*-Cre; *R26R*-H2B- GFP) embryos stained for Prox1, Pdx1, and GFP were digitalized and *xyz*- coordinates for all individual GFP^−^ and GFP^+^ cells in the liver (lv) and pancreato- biliary (pb) buds obtained^46^. Data from individual embryos at E10.5 (a, b; n=8) and E11.0 (c; n=10) were combined and cells plotted by their *xy*- (b) or *yz*-coordinates (a, c). **(d)** Quantification of GFP^+^ cells in LV in genetically labelled embryos (E9.0, n=7; E9.5, n=9; E10.5, n=7; E11.0, n=7) and newborn mice (P1, n=5) shows a constant labelling index of 0.8% of the total liver cell population. No statistically significant differences were detected at the different stages (nonparametric Kruskal-Wallis test, p >0.99). **(e)** Representative IF image of newborn Tg(*Pdx1*-Cre; *R26R*-H2B-GFP) mouse liver. Immunostaining for glutamine synthetase (GS, red) marks hepatocytes near the central vein, whereas CK19 (blue) marks cholangiocytes. Co-staining of GFP with GS (arrows) or CK19 (arrowheads) unveiled Pdx1-derived hepatoblasts capable to differentiate into hepatocytes and cholangiocytes. Scale bar, 100µm. **(f)** Representative IF image of Tg(*Pdx1*-Cre; *R26R*-H2B-GFP) embryos at E9.5. Right panels show boxed region at a higher magnification. Arrowheads indicate GFP^+^genetically labelled cells in PB bud and surrounding LV bud that are positive for CK19. Scale bar, 100µm.

**Extended Data Fig. 6.**
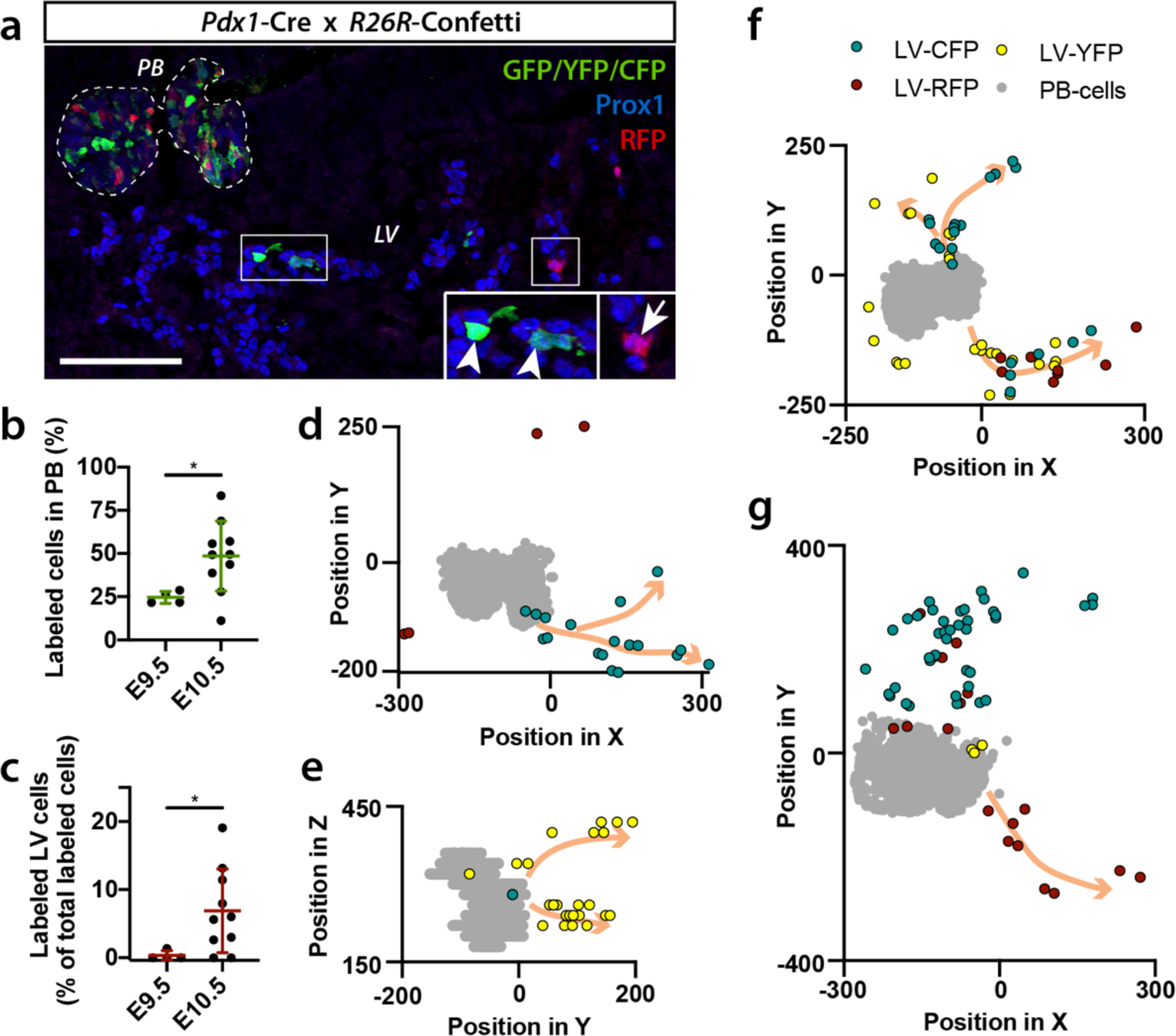
Clonal lineage tracing of pancreato-biliary progenitors shows dispersal of labelled cells in the liver. **(a)** Representative IF image of E10.5 Tg(*Pdx1*-Cre; *R26R*-Confetti) embryos stained with antibodies against Prox1, RFP and GFP. Anti-GFP antibody also detects membrane-bound CFP and cytoplasmic YFP. Dashed white line marks pancreato-biliary (PB) progenitor cells. Insets show higher magnification of the boxed regions highlighting genetically labelled descendants of Pdx1^+^ PB cells expressing YFP (arrowheads) and RFP (arrow) in the liver (LV) bud. Scale bar, 100µm. **(b)** Quantification of labelled PB progenitors in Tg(*Pdx1*-Cre; *R26R*-Confetti) embryos (E9.5, n=4; E10.5, n=10). Genetically labelled cell population increases with developmental stage. Two-tailed Mann-Whitney test, p-value=0.04. **(c)** Quantification of labelled LV progenitors as % of the total labelled cell population in the ventral foregut endoderm of Tg(*Pdx1*-Cre; *R26R*-Confetti) embryos (E9.5, n=4; E10.5, n=10). Two-tailed Mann-Whitney test, p-value=0.03. **(d-g)** Spatial representations of clonal *Pdx1*-Cre lineage tracing experiments in four individual E10.5 Tg(*Pdx1*-Cre; *R26R*-Confetti) embryos. IF data were digitalized and *xyz*-coordinates for individual labelled LV (LV-CFP, LV-RFP, LV-YFP) and PB progenitors (PB-cells) obtained^44^. Cells were plotted according to the *xy*- (d, f, g) or *yz*-coordinates (e). Arrows indicate potential trajectories of labelled cell clusters arising from Pdx1^+^ PB bud and spreading along hepatic chords.

**Extended Data Fig. 7.**
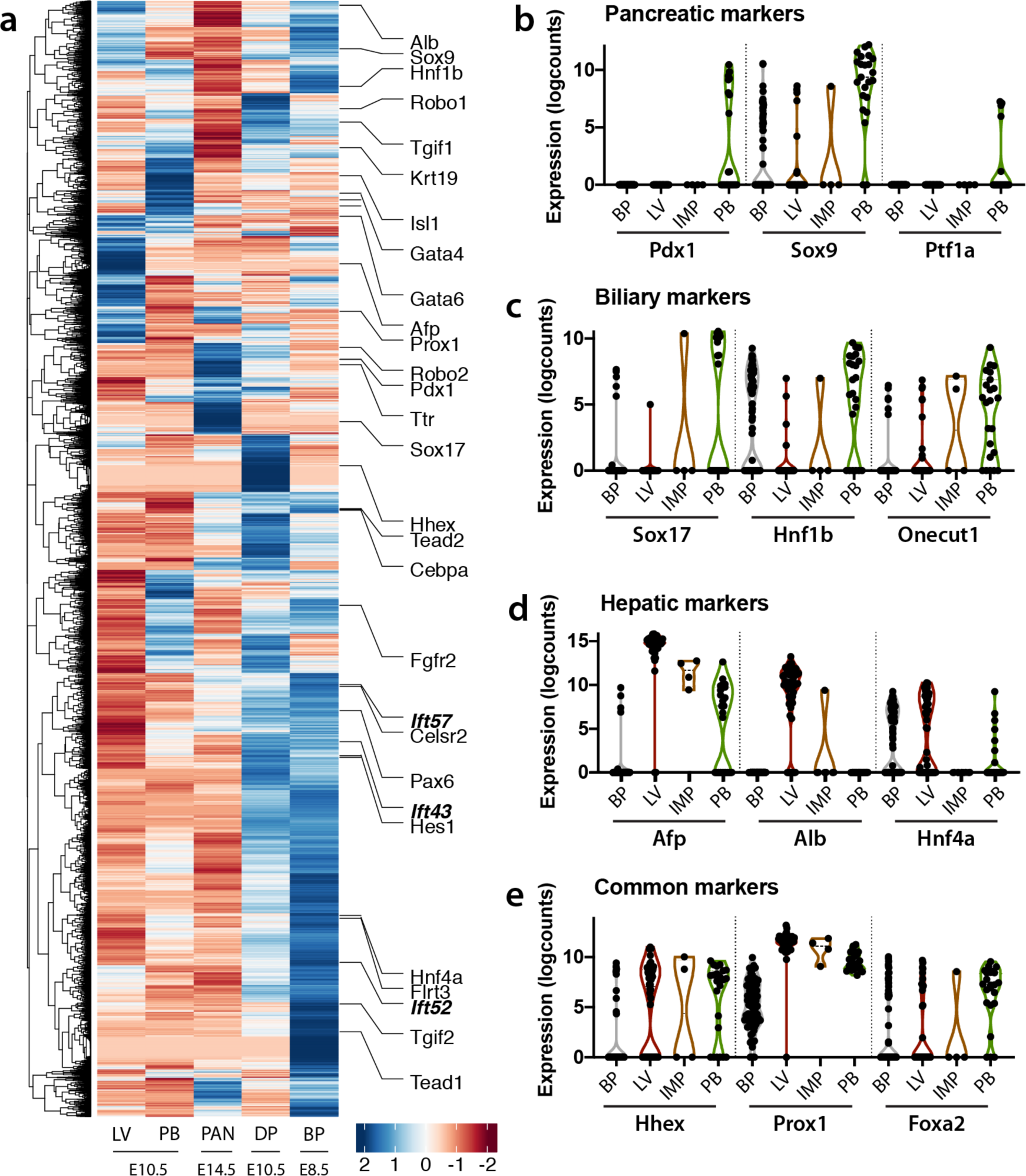
Distinct gene signatures in pancreato-biliary sub- populations. **(a)** Heatmap of sc-RNA-seq dataset from E8.5 bipotent endoderm (BP) progenitors (*a.k.a.* ventral foregut), E10.5 hepatic (LV), dorsal pancreatic (DP), ventral pancreato-biliary cells (PB) and E14.5 pancreatic cells (PAN). Colours represent high (blue) or low (red) normalised log-expression values. Selected genes involved in hepato-pancreatic development are indicated on the right. **(b-e)** Violin plots of normalised log-expression values of selected cell-type-specific genes. IMP, intermediate progenitor cells.

**Extended Data Fig. 8.**
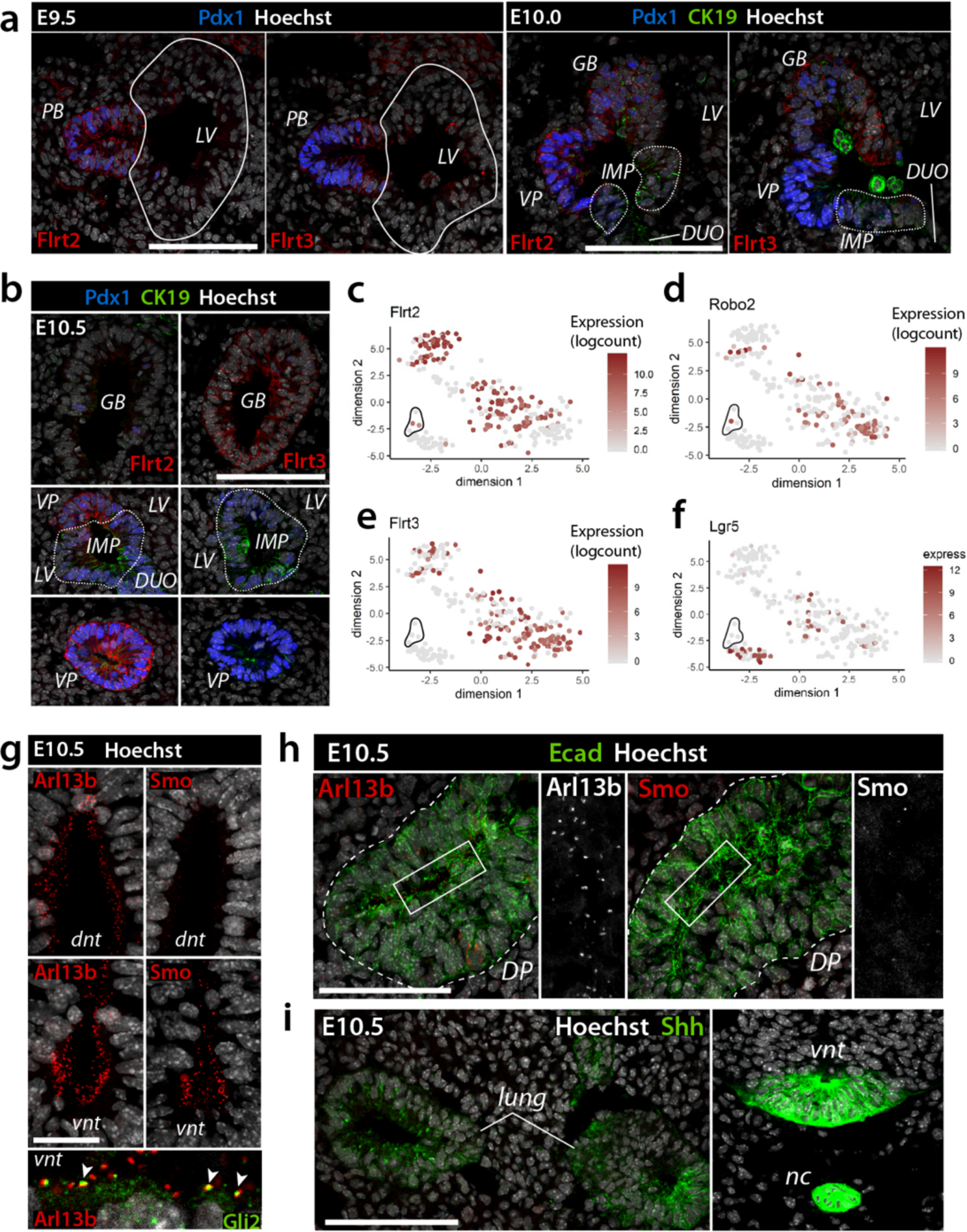
Extended characterization of signalling signatures defining the hepato-pancreato-biliary progenitor domain. **(a,b)** Representative IF images of E9.5-E10.5 embryos stained for the indicated markers. Flrt2 and Flrt3 first co-localize in the same regions of the pancreato-biliary (PB) bud at E9.5, but mark distinct sub-populations within the PB rudiment at later stages. Flrt2 is enriched in the Pdx1^+^ ventral pancreas (VP), while Flrt3 is enriched in the Pdx1^-^ gallbladder (GB). CK19 marks intermediate (IMP) sub-population at the border of duodenum (DUO), liver (LV), and PB bud. Scale bars, 100µm. **(c-f)** t-SNE plots showing the expression of *Flrt2* (c), *Robo2* (d) *Flrt3* (e), and *Lgr5* (f) in the sc-RNA-Seq dataset. Colours span a gradient from red (high expression) to grey (low expression). **(g-i)** Representative IF images of E10.5 embryos stained for cilia markers (Arl13b) and Shh pathway components (Smo, Gli2, Shh). In (g), ventral neural tube (vnt) served as positive control for hedgehog signalling, showing high density of Smo localization at the primary cilia (Arl13b^+^) together with Gli2, while it is absent in dorsal neural tube (dnt)^34, 35^. In (h), IF staining for Arl13b (red) and Smo (red) shows no hedgehog signalling activity in dorsal pancreas, marked by E-cadherin (Ecad; green). In (i), IF staining for Shh (green) shows abundant levels in the lungs, vnt, and notochord (nc), as previously published^34, 35^. Scale bar, 100µm.

**Extended Data Fig. 9.**
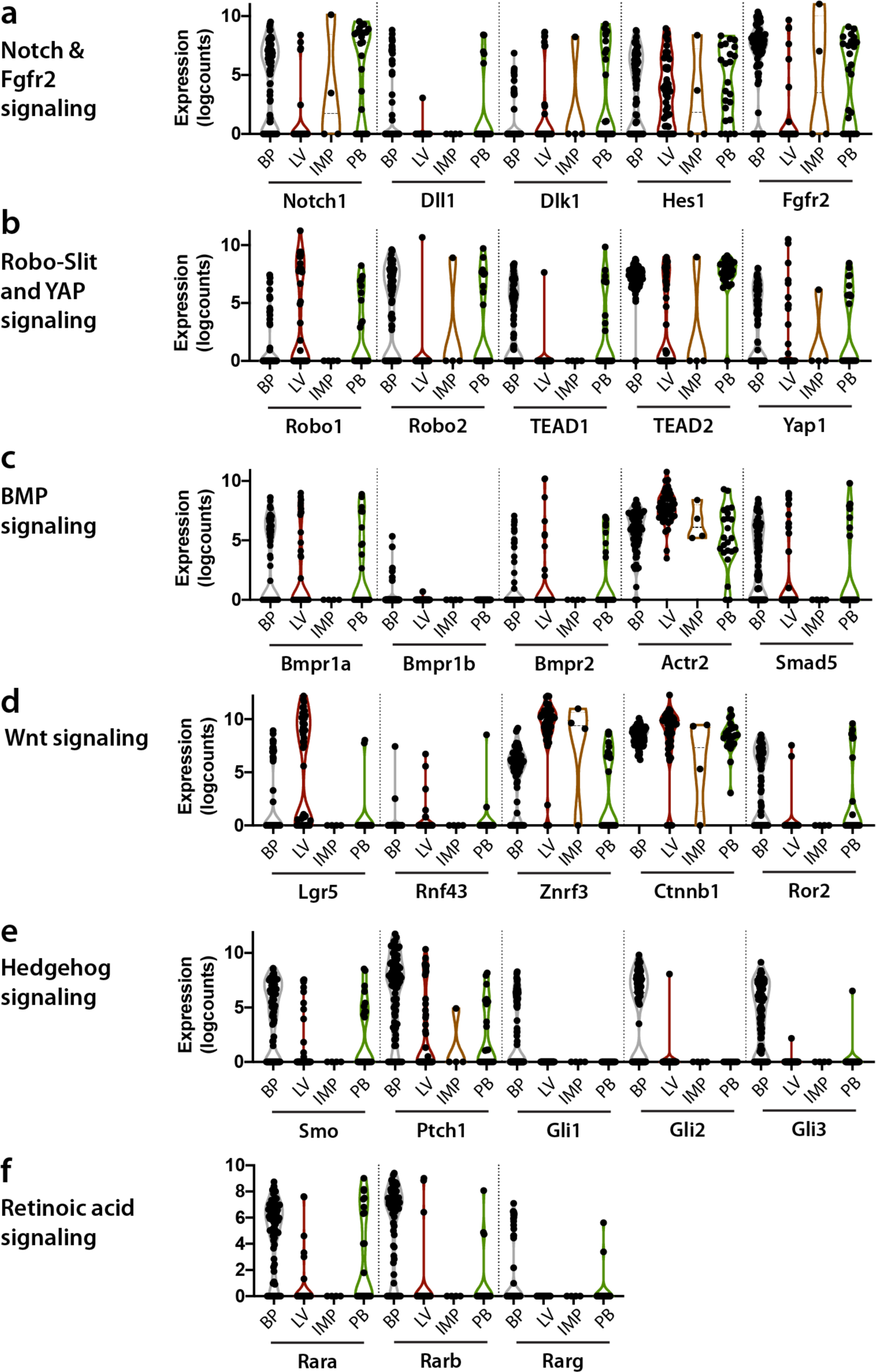
Gene expression signatures of signalling pathways involved in hepato-pancreatic development. **(a-f)** Violin plots of normalised log-expression values of selected components or transducers of Notch and Fgfr2 (a), Robo-Slit and YAP (b), BMP (c), canonical and non-canonical Wnt (d), Hedgehog (e) and Retinoic acid (f) signalling pathways from sc-RNA-seq of E8.5 bipotent endoderm progenitors (BP), E10.5 hepatic (LV), pancreato-biliary cells (PB), and intermediate progenitor (IMP) cells.

## Supplementary Information Note 1: Description of the mathematical models

Several models have been created to describe distinct but similarly plausible hypotheses of the embryonic development of murine liver and ventral pancreas cells. The temporal changes of the cell populations are modelled using systems of ordinary differential equations (ODEs).

### Cell populations of the models

**Table S1.** Description of the cell populations considered in the models, their acronyms and attributed phenotypic markers.

| Acronym | Description | Phenotypic markers |
| --- | --- | --- |
| $BP$ | bipotent progenitor cells | Prox1 |
| $BP_p$ | proliferating bipotent progenitor cells | Prox1 & pH3 |
| $LV$ | hepatic progenitors | Prox1 |
| $LV_p$ | proliferating hepatic progenitors | Prox1 & pH3 |
| $PB$ | pancreato-biliary progenitors | Prox1 & (Pdx1 or Sox17) |
| $PB_p$ | proliferating pancreato-biliary progenitors | Prox1 & (Pdx1 or Sox17) & pH3 |
| $GB$ | gallbladder progenitors | Prox1 & Sox17 |
| $GB_p$ | proliferating gallbladder progenitors | Prox1 & Sox17 & pH3 |
| $VP$ | ventral pancreatic progenitors | Prox1 & Pdx1 |
| $VP_p$ | proliferating ventral pancreatic progenitors | Prox1 & Pdx1 & pH3 |

Rates
| Proliferation processes |  | Differentiation processes |  |
| --- | --- | --- | --- |
| $v_1 = k_1 \cdot [LV]$ | $v_2 = k_2 \cdot [LV_p]$ | $v_{11} = k_{11} \cdot [BP]$ | $v_{13} = k_{13} \cdot [BP]$ |
| $v_3 = k_3 \cdot [PB]$ | $v_4 = k_4 \cdot [PB_p]$ | $v_{17} = k_{17} \cdot [LV]$ | $v_{18} = k_{18} \cdot [PB]$ |
| $v_5 = k_5 \cdot [BP]$ | $v_6 = k_6 \cdot [BP_p]$ | $v_{26} = k_{26} \cdot [PB]$ | $v_{27} = k_{27} \cdot [PB]$ |
| $v_{21} = k_{21} \cdot [GB]$ | $v_{22} = k_{22} \cdot [GB_p]$ | | |
| $v_{23} = k_{23} \cdot [VP]$ | $v_{24} = k_{24} \cdot [VP_p]$ | | |

### Model structures

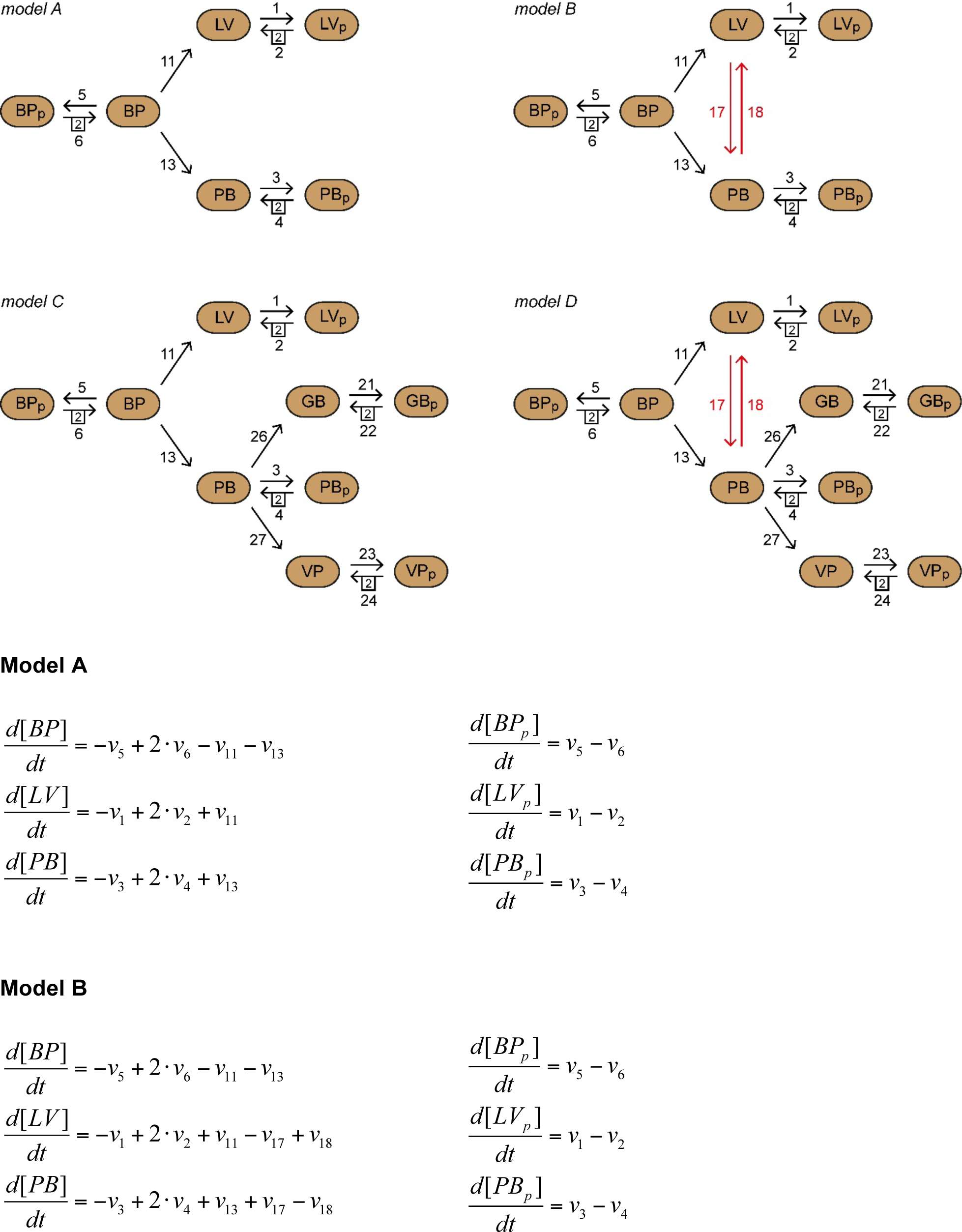
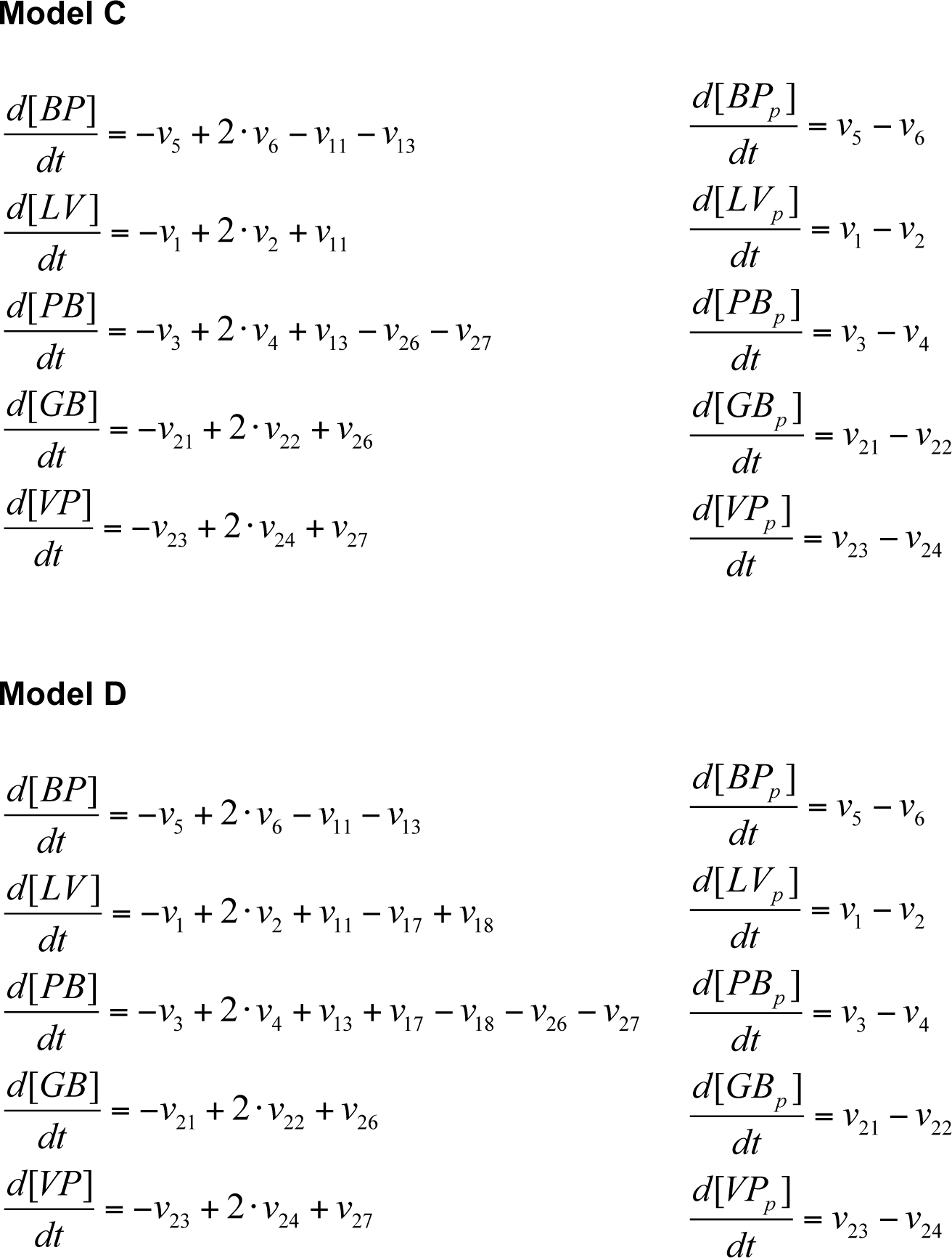

### Parameter estimation

#### Experimental data

Cell counts at different somite stages have been determined as described in the experimental procedures. These counts are listed in the supplemental data file “Supplementary Dataset 1.xlsx”. The cell counts correspond to sums of different cell populations in the models depending on the phenotypic markers attributed to the cell populations (Table S1). For example, the experimentally determined cell count of Prox1-positive cells corresponds to the sum of the calculated cell counts of cell population BP, LV, PB, BP_p_, LV_p_ and PB_p_ in model A (Table S2).

In addition to cell counts, the frequency of proliferating (pH3-positive) cells in foregut cells have been experimentally determined at early time points in development. This frequency corresponds to the ratio between the sum of all pH3-positive cell populations and the sum of all cell populations in the models. The models consider time in hours. We therefore transform somite stage into hours by assuming that a new somite is build every 2h^1, 2^, beginning at *t0 =* 0 with zero somite present.

**Table S2.** Experimental data and their corresponding sums of cell populations in models A and B.

| Column name in ‘Supplementary Dataset 1’ | Models A and B |
| --- | --- |
| Prox1_cell_count | $[BP] + [BP_p] + [LV] + [LV_p] + [PB] + [PB_p]$ |
| Prox1_Pdx1_or_Sox17_cell_count | $[PB] + [PB_p]$ |
| Prox1_pH3_cell_count | $[BP_p] + [LV_p] + [PB_p]$ |
| Prox1_Pdx1_or_Sox17_pH3_cell_count | $[PB_p]$ |
| Frequency of pH3-positive cells<br>(column T / column S) | $\frac{[BP_p] + [LV_p] + [PB_p]}{[BP] + [BP_p] + [LV] + [LV_p] + [PB] + [PB_p]}$ |

**Table S3.** Experimental data and their corresponding sums of cell populations in models C and D.

| Column name in ‘Supplementary Dataset 1’ | Models C and D |
| --- | --- |
| Prox1_cell_count | $[BP] + [BP_p] + [LV] + [LV_p] + [PB] + [PB_p] + [GB] + [GB_p] + [VP] + [VP_p]$ |
| Prox1_Pdx1_or_Sox17_cell_count | $[PB] + [PB_p] + [GB] + [GB_p] + [VP] + [VP_p]$ |
| Prox1_Sox17_cell_count | $[GB] + [GB_p]$ |
| Prox1_Pdx1_cell_count | $[VP] + [VP_p]$ |
| Prox1_pH3_cell_count | $[BP_p] + [LV_p] + [PB_p] + [GB_p] + [VP_p]$ |
| Prox1_Pdx1_or_Sox17_pH3_cell_count | $[PB_p] + [GB_p] + [VP_p]$ |
| Prox1_Sox17_pH3_cell_count | $[GB_p]$ |
| Prox1_Pdx1_pH3_cell_count | $[VP_p]$ |
| Frequency of pH3-positive cells<br>(column T / column S) | $\frac{[BP_p] + [LV_p] + [PB_p] + [GB_p] + [VP_p]}{[BP] + [BP_p] + [LV] + [LV_p] + [PB] + [PB_p] + [GB] + [GB_p] + [VP] + [VP_p]}$ |

#### Optimization method

We determined the parameters of each model, i.e. all corresponding rate parameters (*k_i_*) and initial cell counts of the cell populations at *t_0_ =* 0, that maximise the likelihood function given the experimental data. To do so, we used a deterministic optimization algorithm implemented in the D2D Toolbox for Matlab^3, 4^ and sampled for each model 1000 initial parameter sets with Latin hypercube sampling. Rate parameters were limited to the range from 0.001 to 1 with the exception of *k_2_*, which ranged from 0.001 to 1000. Initial conditions were limited to the range from 0.5 to 400. We chose the parameter set of the best fit out of the 1000 optimization runs. The chosen parameter set of each model is listed in Table S4 and Table S5.

#### Parameters of the models

**Table S4.** Rate parameters of the models [1/h].

| Model | A | B | C | D |
| --- | --- | --- | --- | --- |
| $k_1$ | 0.069 | 0.073 | 0.070 | 0.064 |
| $k_2$ | 1.89 | 1.36 | 2.05 | 1.48 |
| $k_3$ | 0.041 | 0.028 | 0.0021 | 0.0055 |
| $k_4$ | 0.33 | 0.21 | 0.12 | 0.18 |
| $k_5$ | 0.12 | 0.001 | 0.001 | 0.020 |
| $k_6$ | 0.023 | 0.035 | 0.022 | 0.035 |
| $k_{11}$ | 0.34 | 0.001 | 0.44 | 0.001 |
| $k_{13}$ | 0.38 | 0.15 | 1 | 1 |
| $k_{17}$ | | 0.013 | | 0.001 |
| $k_{18}$ | | 0.057 | | 0.040 |
| $k_{21}$ | | | 0.080 | 0.083 |
| $k_{22}$ | | | 0.26 | 0.29 |
| $k_{23}$ | | | 0.073 | 0.077 |
| $k_{24}$ | | | 0.19 | 0.22 |
| $k_{26}$ | | | 0.0056 | 0.0041 |
| $k_{27}$ | | | 0.0050 | 0.0036 |

**Table S5.** Initial conditions of the models [cells].

| Model | A | B | C | D |
| --- | --- | --- | --- | --- |
| $BP$ | 97 | 125 | 124 | 135 |
| $BP_p$ | 88 | 119 | 113 | 126 |
| $LV$ | 0 | 0 | 0 | 0 |
| $LV_p$ | 0 | 0 | 0 | 0 |
| $PB$ | 0 | 0 | 0 | 0 |
| $PB_p$ | 0 | 0 | 0 | 0 |
| $GB$ | 0 | 0 | 0 | 0 |
| $GB_p$ | 0 | 0 | 0 | 0 |
| $VP$ | 0 | 0 | 0 | 0 |
| $VP_p$ | 0 | 0 | 0 | 0 |

#### Model comparison

We compared models using the relative likelihood to identify the model that explains the data best. The relative likelihood *ω* of model M2 with respect to model M1 is defined as:

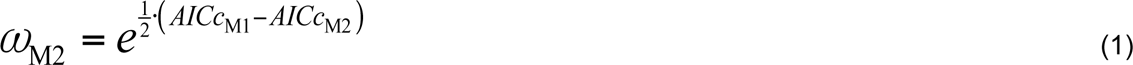

where AICc stands for the corrected Akaike information criterion (AICc)^5, 6^. The AICc was calculated by:

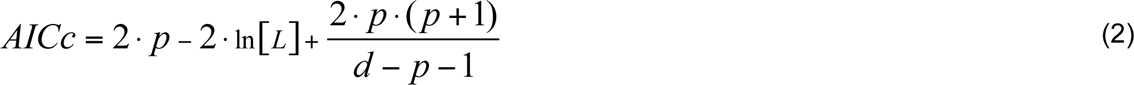

where *p* is the number of parameters of the tested model, *d* denotes the number of data points, and ln[*L*] is the log-likelihood of the model parameter values given the data. Calculations were done with Matlab (R2017b, The MathWorks Inc., Natick, MA).

#### Numerical simulations

Simulations were done with Mathematica 11.0 (Wolfram Research).

## Supplementary Information Note 2: Generation of the Tg(*Prox1*-rtTA) transgenic mouse line

The Tg(*Prox1*-rtTA) transgenic mouse line was generated by modifying a mouse bacterial artificial chromosome (BAC) clone (RP23-360I16) spanning the Prox1 gene and its up- and downstream regulatory sequences. The transgenic expression cassette encodes for a reverse-tetracycline controlled transactivator (rtTA) fused to mCherry via a 2A peptide. The 2A peptide possesses a self-cleaving activity and allows separation of rtTA and mCherry proteins after translation. The rtTA-2A- mCherry cassette was then inserted through homologous recombination into the RP23-360I16 BAC clone, which contains an approximately 196-kb-long mouse genomic contig, spanning from approximately 115 kb upstream of the *Prox1* start codon to approximately 81 kb downstream of the start codon and harbours all regulatory elements that have been proposed to regulate the *Prox1* expression^7^.

Briefly, two sequences (called Box A and Box B) homologous to the genomic region around the *Prox1* start codon, lying in exon 2 of the *Prox1* gene, were cloned upstream and downstream of the rtTA-2A-mCherry cassette into the pLD53.SCAEB shuttle vector^8^. Box A corresponded to a 603 bp-long region of the *Prox1* gene, 31 bp-upstream of the start codon, while Box B corresponded to a 507 bp-long region of the *Prox1* gene starting 43 bp-downstream of the start codon. The rtTA-2A-mCherry cassette was introduced into the second exon of the *Prox1* gene via a two-step homologous recombination process, as previously reported^8^. Successful integration of the rtTA-2A-mCherry cassette into the BAC clone was confirmed by PCR and Southern Blotting. Next, BAC DNA was purified and introduced into embryos of B6D2F2 hybrid mice by pronuclear injection at the MDC Transgenic Facility. Injected embryos were transplanted into pseudo-pregnant females and genetic status of offspring was determined by PCR.

**Table S6.**
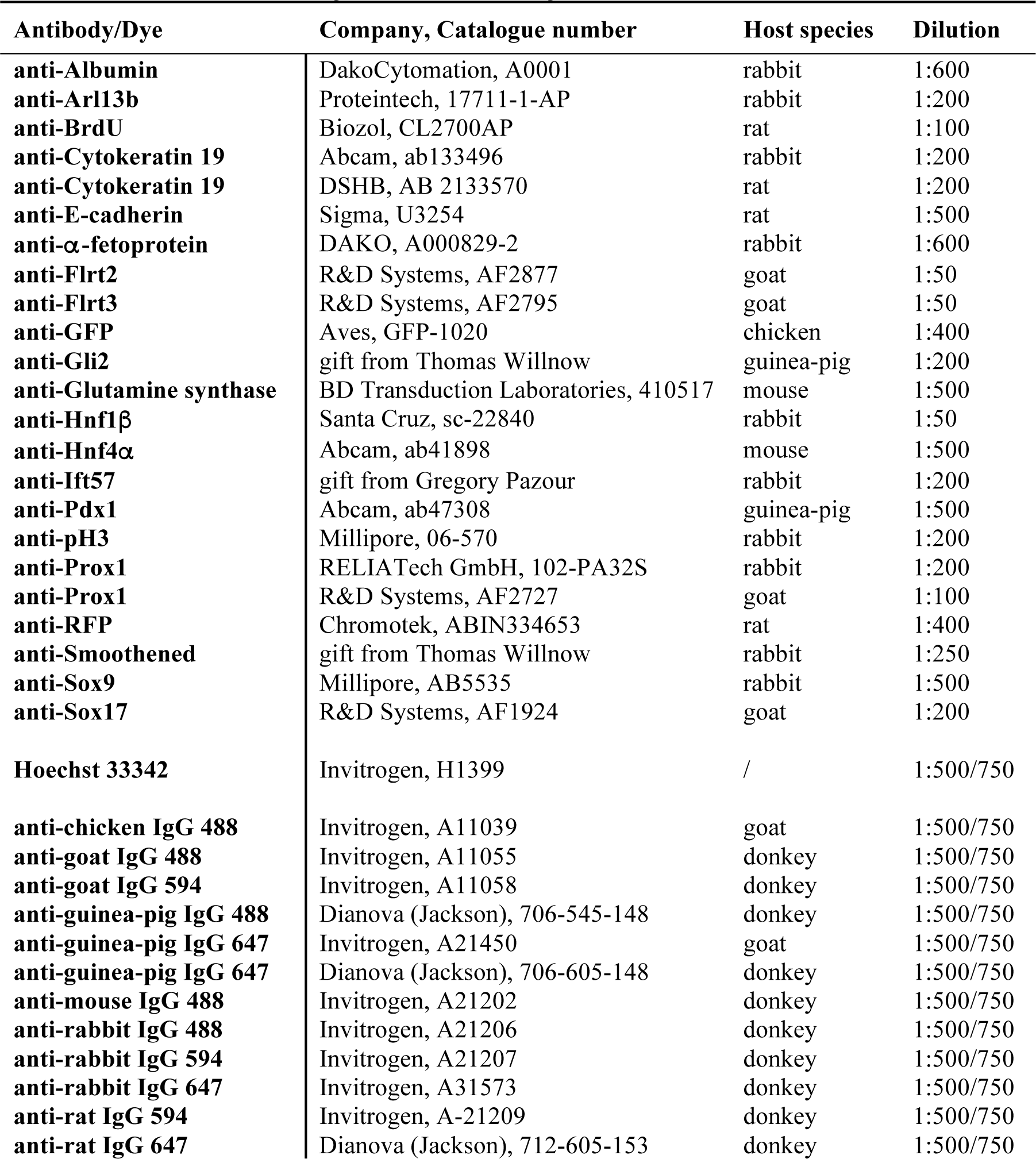
List of Primary and Secondary Antibodies

| Antibody/Dye | Company, Catalogue number | Host species | Dilution |
| --- | --- | --- | --- |
| anti-Albumin | DakoCytomation, A0001 | rabbit | 1:600 |
| anti-Arl13b | Proteintech, 17711-1-AP | rabbit | 1:200 |
| anti-BrdU | Biozol, CL2700AP | rat | 1:100 |
| anti-Cytokeratin 19 | Abcam, ab133496 | rabbit | 1:200 |
| anti-Cytokeratin 19 | DSHB, AB 2133570 | rat | 1:200 |
| anti-E-cadherin | Sigma, U3254 | rat | 1:500 |
| anti-α-fetoprotein | DAKO, A000829-2 | rabbit | 1:600 |
| anti-Flrt2 | R&D Systems, AF2877 | goat | 1:50 |
| anti-Flrt3 | R&D Systems, AF2795 | goat | 1:50 |
| anti-GFP | Aves, GFP-1020 | chicken | 1:400 |
| anti-Gli2 | gift from Thomas Willnow | guinea-pig | 1:200 |
| anti-Glutamine synthase | BD Transduction Laboratories, 410517 | mouse | 1:500 |
| anti-Hnf1β | Santa Cruz, sc-22840 | rabbit | 1:50 |
| anti-Hnf4α | Abcam, ab41898 | mouse | 1:500 |
| anti-Ift57 | gift from Gregory Pazour | rabbit | 1:200 |
| anti-Pdx1 | Abcam, ab47308 | guinea-pig | 1:500 |
| anti-pH3 | Millipore, 06-570 | rabbit | 1:200 |
| anti-Prox1 | RELIAtech GmbH, 102-PA32S | rabbit | 1:200 |
| anti-Prox1 | R&D Systems, AF2727 | goat | 1:100 |
| anti-RFP | Chromotek, ABIN334653 | rat | 1:400 |
| anti-Smoothened | gift from Thomas Willnow | rabbit | 1:250 |
| anti-Sox9 | Millipore, AB5535 | rabbit | 1:500 |
| anti-Sox17 | R&D Systems, AF1924 | goat | 1:200 |
| **Hoechst 33342** | Invitrogen, H1399 | / | 1:500/750 |
| **anti-chicken IgG 488** | Invitrogen, A11039 | goat | 1:500/750 |
| **anti-goat IgG 488** | Invitrogen, A11055 | donkey | 1:500/750 |
| **anti-goat IgG 594** | Invitrogen, A11058 | donkey | 1:500/750 |
| **anti-guinea-pig IgG 488** | Dianova (Jackson), 706-545-148 | donkey | 1:500/750 |
| **anti-guinea-pig IgG 647** | Invitrogen, A21450 | goat | 1:500/750 |
| **anti-guinea-pig IgG 647** | Dianova (Jackson), 706-605-148 | donkey | 1:500/750 |
| **anti-mouse IgG 488** | Invitrogen, A21202 | donkey | 1:500/750 |
| **anti-rabbit IgG 488** | Invitrogen, A21206 | donkey | 1:500/750 |
| **anti-rabbit IgG 594** | Invitrogen, A21207 | donkey | 1:500/750 |
| **anti-rabbit IgG 647** | Invitrogen, A31573 | donkey | 1:500/750 |
| **anti-rat IgG 594** | Invitrogen, A-21209 | donkey | 1:500/750 |
| **anti-rat IgG 647** | Dianova (Jackson), 712-605-153 | donkey | 1:500/750 |

